# Biophysical modeling of frontocentral ERP generation links circuit-level mechanisms of action-stopping to a behavioral race model

**DOI:** 10.1101/2023.10.25.564020

**Authors:** Darcy A. Diesburg, Jan R. Wessel, Stephanie R. Jones

**Affiliations:** Department of Neuroscience, Brown University, Providence, RI, USA; Department of Psychological and Brain Sciences, University of Iowa, Iowa City, IA, USA; Department of Neurology, Carver College of Medicine, University of Iowa Hospitals and Clinics, Iowa City, IA, USA; Center for Neurorestoration and Neurotechnology, Providence VA Medical Center, RI, USA

## Abstract

Human frontocentral event-related potentials (FC-ERPs) are ubiquitous neural correlates of cognition and control, but their generating multiscale mechanisms remain mostly unknown. We used the Human Neocortical Neurosolver(HNN)’s biophysical model of a canonical neocortical circuit under exogenous thalamic and cortical drive to simulate the cell and circuit mechanisms underpinning the P2, N2, and P3 features of the FC-ERP observed after Stop-Signals in the Stop-Signal task (SST). We demonstrate that a sequence of simulated external thalamocortical and cortico-cortical drives can produce the FC-ERP, similar to what has been shown for primary sensory cortices. We used this model of the FC-ERP to examine likely circuit-mechanisms underlying FC-ERP features that distinguish between successful and failed action-stopping. We also tested their adherence to the predictions of the horse-race model of the SST, with specific hypotheses motivated by theoretical links between the P3 and Stop process. These simulations revealed that a difference in P3 onset between successful and failed Stops is most likely due to a later arrival of thalamocortical drive in failed Stops, rather than, for example, a difference in effective strength of the input. In contrast, the same model predicted that early thalamocortical drives underpinning the P2 and N2 differed in both strength and timing across stopping accuracy conditions. Overall, this model generates novel testable predictions of the thalamocortical dynamics underlying FC-ERP generation during action-stopping. Moreover, it provides a detailed cellular and circuit-level interpretation that supports links between these macroscale signatures and predictions of the behavioral race model.

**Significance statement:** The frontocentral event-related potential (FC-ERP) is an easily-measurable neural correlate of cognition and control. However, the cortical dynamics that produce this signature in humans are complex, limiting the ability of researchers to make predictions about its underlying mechanisms. In this study, we used the biophysical model included in the open-source Human Neocortical Neurosolver software to simulate and evaluate the likely cellular and circuit mechanisms that underlie the FC-ERP in the Stop-Signal task. We modeled mechanisms of the FC-ERP during successful and unsuccessful stopping, generating testable predictions regarding Stop-associated computations in human frontal cortex. Moreover, the resulting model parameters provide a starting point for simulating mechanisms of the FC-ERP and other frontal scalp EEG signatures in other task conditions and contexts.

## Introduction

Frontocentral event-related potentials (FC-ERPs) from human electroencephalography (EEG) are ubiquitous proxy signatures of higher-order cognition and control (e.g., Holroyd & Coles, 2002; Falkenstein et al., 2000; Hajcak et al., 2006; Polich, 2007; Proudfit, 2015). For example, they index inhibitory control processes in the Stop-Signal Task (SST; de Jong et al., 1990; Kok et al., 2004). In the SST, participants respond to imperative stimuli and try to cancel ongoing responses when presented with an infrequent “Stop-Signal’’ on a subset of trials (Logan, 1994; Logan & Cowan, 1984a; Verbruggen & Logan, 2009). The Stop-Signal-aligned FC-ERP contains a complex of peaks within approximately 300ms (e.g., the P2, N2, and P3) associated with inhibitory performance. A large, positive-going P3 is observed in Stop but not Go trials (de Jong et al., 1990; Enriquez-Geppert et al., 2010; Huster et al., 2013; Kok et al., 2004), onsets earlier in successful than failed Stop trials (Wessel & Aron, 2015), and has an onset latency that correlates with Stop-Signal reaction time (SSRT, an estimation of how long stopping requires; Huster et al., 2020; Wessel & Aron, 2015). The preceding, negative-going N2’s onset and amplitude also vary with stopping success (Huster et al., 2010, 2011, 2013; Ramautar et al., 2004), though the direction of these effects are mixed (c.f., Dimoska et al., 2006; Senderecka, 2016; Senderecka et al., 2012).

Despite theories associating FC-ERP waveform changes with stopping, the detailed multiscale neural mechanisms generating the FC-ERP and its condition-dependent changes have not been delineated. In this study, we used Human Neocortical Neurosolver (HNN), whose foundation is a biophysical model of the canonical neocortical column under thalamocortical and cortico-cortical drive, to predict multiscale cell and circuit dynamics producing the FC-ERP in the SST. We specifically tested theorized links between these neural mechanisms and predictions of the well-known “horse-race” model, a computational model of behavior in the SST. This model assumes the outcome of Stop trials is determined by the winner of a race between pro-kinetic Go and anti-kinetic Stop processes (Logan & Cowan, 1984a, 1984b). According to the logic of this horse-race model, any *neural* signature of the Stop process that starts earlier should lead to more successful stopping. Consequently, the onset of the Stop-Signal-aligned frontocentral P3 has been proposed to index the timing of this Stop process, since it onsets earlier on successful than failed Stop trials (e.g., Wessel & Aron, 2015). However, though a race model conceptualization aligns with canonical characteristics of the grand-average P3, it does not necessarily follow that underlying neural mechanisms support this interpretation. ERPs are generated by complex, overlapping laminar dynamics (e.g., Jones, 2015; Jones et al., 2007; Lindén et al., 2011; Neymotin et al., 2020; Reimann et al., 2013). Based on the averaged FC-ERP alone, it is impossible to know whether the observed change in onset timing between successful and failed Stop trials is due to an earlier onset of the underlying neural mechanism (i.e., those purportedly underlying stopping), or due to a stronger effective strength of the same mechanism (which would also lead to an earlier emergence of a significant FC-ERP feature).

HNN was explicitly designed to simulate the multiscale biophysical origin of human M/EEG signals. HNN accounts for both local circuit activity and exogenous driving influences from subcortical thalamic and higher/lower order cortical areas (Neymotin et al., 2020). As such, it provides a unique opportunity to predict whether the likely neural dynamics underlying condition differences in the FC-ERP support or negate the race model interpretation of P3. We applied HNN for the first time to signals from frontal cortex to model the mechanisms underlying the FC-ERP during successful and failed Stop trials in the SST.

## Methods

FC-ERPs used to fit HNN models were extracted from an open-source EEG dataset collected during a SST (https://osf.io/v3a78/). The data collection procedures are described in full in Wessel (2020) and briefly here. Findings from this dataset have been published previously in whole or in part in other investigations that have not included biophysical modeling (Dutra et al., 2018; Dykstra et al., 2020; Soh & Wessel, 2021; Waller et al., 2021; Wessel, 2018a, 2020, p. 202; Wessel & Huber, 2019). All code for analyses specific to this study can be found on Github at https://github.com/darcywaller/HNN_StopSignal_FCERP.

### Participants

Participants were 234 healthy young adults (mean age: 22.7, SEM: 0.43, 137 female, 25 left-handed) who participated in research studies at the University of Iowa (IRB #201511709). Participants were compensated for participation monetarily or with course credit.

### Behavioral paradigm

All participants performed a computerized SST (on Linux using Psychtoolbox; Brainard, 1997) during simultaneous scalp EEG recording (see **Figure 2A**). Individuals were told to respond to black arrows (Go signals) appearing on a gray background as quickly as possible using ‘q’ and ‘p’ keys on a keyboard for “left” and “right”, respectively. On a third of trials, following a delay, participants received a second, infrequent Stop-Signal and saw the arrow turn red, cueing them to halt their response if possible. Initial Stop-Signal delay (SSD) was 250ms and was adjusted throughout the task (decreased by 50ms for failed Stops and increased by 50ms for successful Stops). Participants were instructed to equally prioritize responding quickly and stopping when possible. Feedback was given when necessary by researchers during the block breaks to ensure adherence to task rules.

### EEG data collection

EEG data were collected using either the Brain Products actiChamp or passive MR plus cap. Both caps contained 62 electrodes. The passive cap was used with two additional electrodes placed on the left canthus and the orbital bone below the left eye. The Fz and Pz electrodes were used as ground and reference contacts, respectively. Data were digitized at a rate of 500Hz, recording filters set at a highpass of 10s and lowpass of 1000Hz, and impedance was kept below 10mΩ.

### Preprocessing

This analysis utilized the already-preprocessed datasets provided in the OSF upload from Wessel (2020). We described the preprocessing pipeline here briefly, but see Wessel (2020) and associated code at https://osf.io/v3a78/ for complete details. Data were preprocessed using custom MATLAB code using the EEGLAB toolbox (Delorme & Makeig, 2004). Once imported into EEGLAB in MATLAB, the data were filtered with a high-pass FIR filter at 0.3Hz and a low-pass FIR filter at 30Hz. The data were segmented into 1s epochs and screened for periods in which amplitude or kurtosis exceeded five times the mean. Any segment in which this threshold was met was considered to contain an artifact and was removed. Following cleaning, data were re-referenced to the common average. A temporal infomax ICA decomposition algorithm (Bell & Sejnowski, 1995) with extension to sub-Gaussian sources (Lee et al., 1999) was performed and the resulting component matrix was algorithmically screened for components representing eye movement and electrode artifacts using outlier statistics (Dutra et al., 2018; Waller et al., 2021; Wessel, 2018a). Independent components identified as artifacts were subtracted from the data.

### Identification of independent component contributing to P3 latency difference in FC-ERP

Prior work on the neural generators of the FC-ERP during the SST implicates generators for the N2/P3 in medial prefrontal cortex (mPFC; specifically, encompassing the presupplementary motor area and midcingulate cortex; preSMA / MCC; Huster et al., 2011). Here, we did not compute an EEG inverse solution because we did not have access to participants’ structural MRIs. We chose not to use a template brain for source reconstruction because source-localization of mPFC dipoles is imprecise in the absence of individual structural scans because of the impacts of individual variation in cingulate morphology on the N2/P3 complex (Huster et al., 2014). Rather, we constrained our data in a manner that maximized our chances of studying activity from *frontal medial generators* and allowed us to address *a priori* hypotheses about the P3. Specifically, we used ICA to isolate the independent component (IC) in our data that accounts for the successful versus failed Stop condition difference in the onset of the P3 in the SST (Wessel, 2018b), and examined the contribution from this component to time-series voltage at channel FCz. ICA applied across time yields low-dimensional spatial components of the data that are predicted to come from independent underlying neural generators, though this analysis is agnostic to underlying anatomy. In each subject, we identified the independent component 1) whose channel weights were maximal at frontocentral electrodes Fz, F1, F2, FCz, FC1, FC2, Cz, C1, or C2, and 2) whose reconstructed (in channel space) time-series data related to the P3 onset difference in the all-IC data most strongly. This second condition was assessed by calculating the difference between the failed and successful Stop ERP in the time range of 200-500ms post-Stop-Signal in both the candidate-IC and all-IC data, then correlating the two. Once the component with the correct topography and strongest correlation with P3 onset difference was identified, we then removed all other ICs from the data and reconstructed our time-series at each channel using the activation of this single independent component. This procedure has been used successfully in previous work to isolate the IC (and assumed underlying independent neural generator) that accounts for the P3 onset difference (Dutra et al., 2018; Dykstra et al., 2020; Waller et al., 2021; Wessel & Huber, 2019), and was used here with the motivation of isolating the neural generator of activity associated most closely with the Stop process. It is worth noting that though the algorithmic approach targets large-amplitude condition differences in the P3 time frame, the resulting IC also includes P2 and N2 features when ERPs are extracted (see **Figure 2C**).

### Event-related potentials

From cleaned data reconstructed using the independent component described above, we extracted ERPs at electrode FCz (central channel from the frontocentral ROI in Wessel & Aron, 2015; Skippen et al., 2020) by epoching the data from 100ms preceding to 500ms following the Stop-Signal for failed and successful Stop trials. In addition, matched Go trial ERPs were made by identifying Go trials that followed each Stop trial as closely as possible (and therefore contained the same SSD in the staircase). These Go trials were epoched around the time of SSD, even though no Stop-Signal occurred on those trials, yielding a control time period in which no Stop-related processes should be active. Condition grand averages were made by averaging over all trials and subjects within conditions. All ERPs were baseline-corrected with the mean-subtraction method using a period of 100ms preceding the Stop-Signal (or the SSD, in the case of matched Go trials).

### Quantifying ERP onsets and peaks

The timing of average P2, N2, and P3 peak amplitudes were determined for each participant by taking the maximum of the subject-average ERP in the 100-200ms post-Stop-Signal time period, the minimum 150-250ms post-Stop-Signal, and the maximum during the whole-ERP window, respectively. N2 peak latency was quantified as the timing of the same minimum in the subject-average ERP 150-250ms post-Stop-Signal. P3 onset latency was quantified as in Wessel and Aron (2015). For each Stop condition (successful and failed) within-subjects, a pool of single-trial ERPs from all Stop trials was tested against the pool of matched Go trials for that Stop success condition using bootstrapped Monte-Carlo *t*-tests with 1,000 permutations. The resulting vector of *p* values across time-points was corrected for multiple comparisons using the Benjamini-Hochberg family-wise error-rate correction (Benjamini et al., 2006) to *p*<0.05. Taking the corrected *p*-value vector, we identified the first time point prior to the P3 peak which contained a significant difference between Stop and Go trial amplitudes to identify the first time point at which the ERPs for Stop and Go conditions significantly diverged during the P3’s onset.

### Human Neocortical Neurosolver (HNN)

*Overview.* HNN is a neural modeling software whose foundation is a biophysically principled neocortical column model under exogenous thalamocortical and cortico-cortical synaptic drive (**Figure 1**). It is explicitly designed to simulate the primary current dipoles that generate EEG and MEG signals based on their biophysical origin from current flow in pyramidal neuron dendrites, and with enough detail to connect to microcircuit dynamics. A comprehensive description of this model and empirical support for its components can be found in Neymotin et al. (2020; also see **Figure 1**). The software and tutorials for use are distributed at https://hnn.brown.edu and all code is available on Github. Models in this investigation were run with the 1.3.2 release of HNN GUI in combination with the updated Layer 5 calcium dynamics file developed by Kohl et al. (2022).

**Figure 1.**
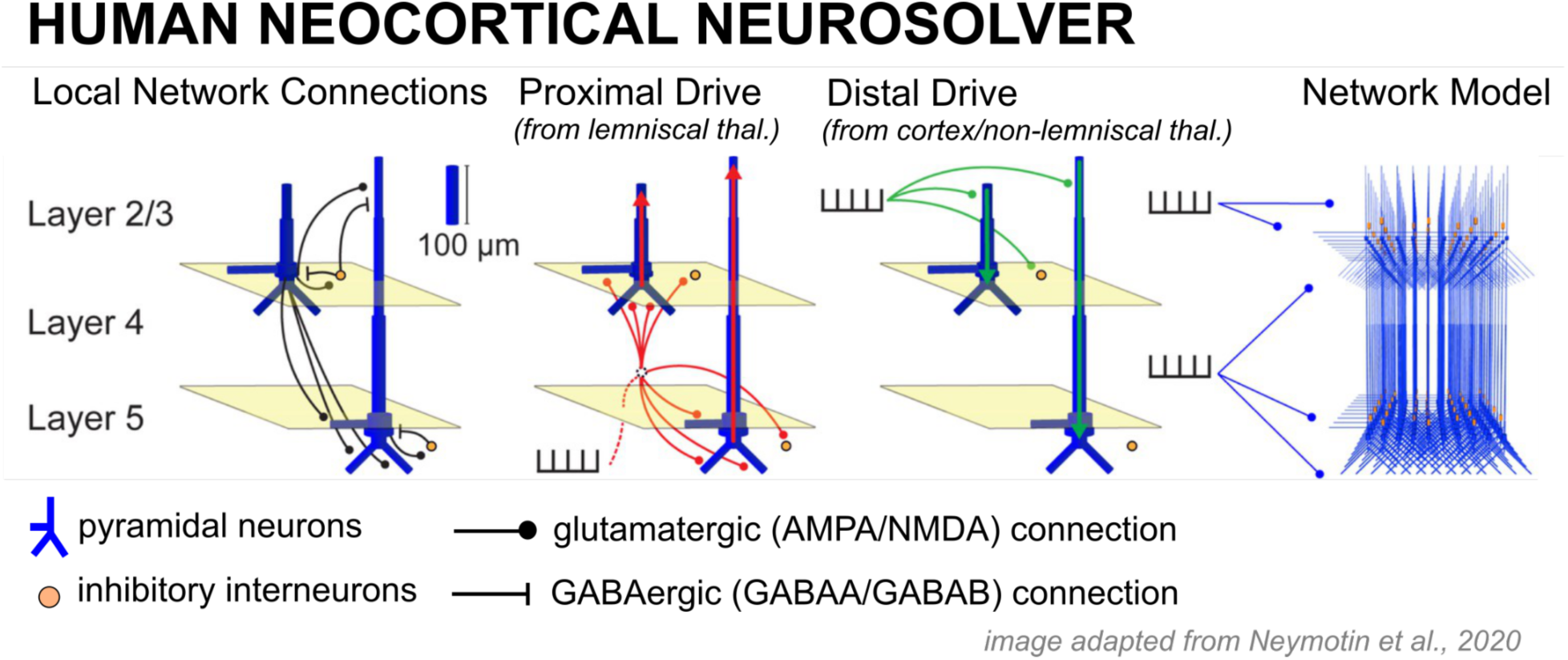
Diagram of the Human Neocortical Neurosolver (HNN) model of a canonical neocortical column. Activity in a laminar network of excitatory pyramidal and inhibitory basket cells are driven by the delivery of spikes or trains of spikes in the form of proximal drive to an abstracted Layer 4 or distal drive to Layers 2/3.

**Figure 2.**
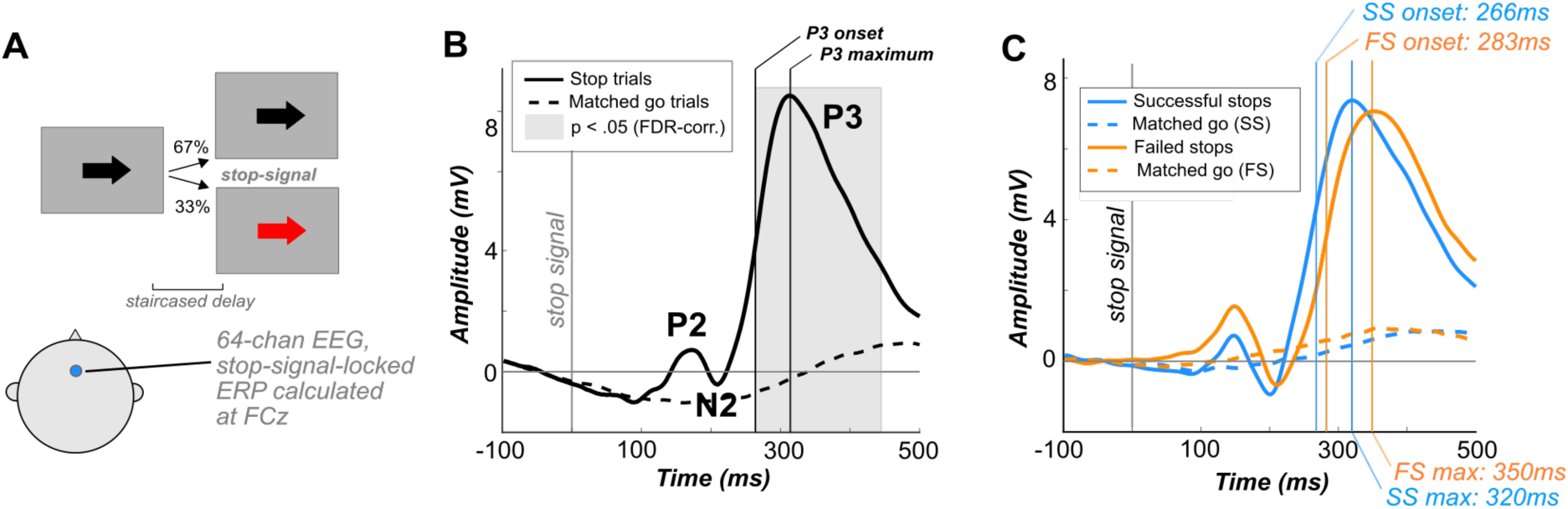
The Stop-Signal Task (SST) and Stop-Signal locked FC-ERPs. A) Participants performed a visual SST while EEG was recorded. FC-ERPs were extracted from electrode FCz. B) P3 onset was calculated using the procedure described in Wessel and Aron (2015), in which permutation testing was applied within-subjects to identify the time point when the Stop trial ERPs significantly deflected from Go trial ERPs. C) Grand-average Stop-Signal locked FC-ERPs in our sample. The onset and peak latency of the P3 and peak latency of the N2 were earlier in successful compared to failed Stops.

*Neocortical column model.* The model consists of individual cells arranged in a laminar structure that comprises a canonical neocortical column. The network contains multi-compartment pyramidal cells and inhibitory neurons simulated as point neurons in Layers 2/3 and Layer 5 of cortex. Layer 4 is not explicitly modeled but assumed to directly relay feedforward input from the thalamus to cells in the other layers, effectively contacting the proximal dendrites of the pyramidal neurons (see Proximal drive in **Figure 1**). Cells are arranged in an equidistant grid with a 3:1 pyramidal-interneuron ratio, with the default model size of 100 pyramidal cells in each layer.

In order to establish a HNN model for medial frontal cortex, we made several changes to the default values in the local neocortical column model (previously developed to account for data from primary sensory cortex). The literature on receptor concentration within layers of medial frontal cortex (e.g., including midcingulate cortex) suggests that this area of the cortex differs from other regions in several ways. In particular, there is an abundance of AMPA-facilitated interconnections in pyramidal cells, resulting in networks that are “hyper-reciprocally connected” (Wang et al., 2006). There is high NMDA and GABAb binding across layers and high GABAa binding in layers I-III specifically (Palomero-Gallagher et al., 2009; Vogt, 2016). These properties are proposed to factor into the high propensity of medial prefrontal neurons for augmentation and potentiation following transient input (e.g., Hempel et al., 2000). In line with these observations, we made several changes to the local connectivity of the model underlying HNN: 1) increased NMDA weights from layer 2 pyramidal cells to layer 2 pyramidal cells by 50% and from layer 5 pyramidal cells to layer 5 pyramidal cells by 50%; 2) increased AMPA weights from layer 2 pyramidal cells to layer 2 pyramidal cells by 50%, from layer 2 pyramidal to layer 5 pyramidal cells by 100%, and from layer 5 pyramidal cells to layer 5 pyramidal cells by 50%; 3) increased GABAa weights from layer 2 basket cells to layer 2 pyramidal cells by 100%; and 4) increased GABAb weights from layer 2 basket cells to layer 2 pyramidal cells by 100% and from layer 5 basket to layer 5 pyramidal cells by 200%. All other local network parameters remained fixed to default values determined by prior work (See **Figure 3-1** in the Extended Data). As described below, we adjusted parameters representing the exogenous drive to the local network to test hypotheses on the generation of the FC-ERP.

**Figure 3.**
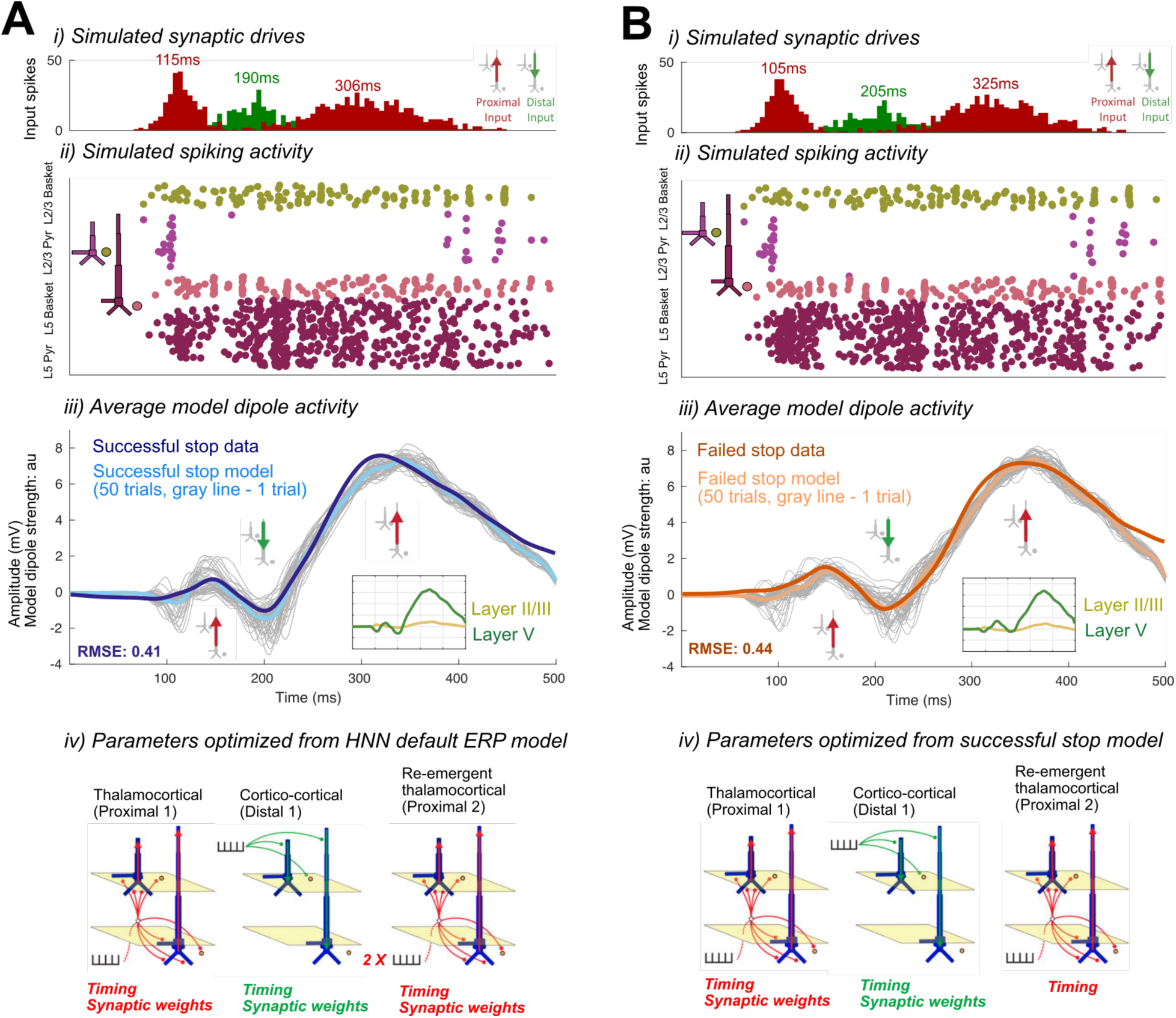
Optimized HNN models for the successful Stop (A) and failed Stop FC-ERP (B). i) All-trials histograms of thalamocortical and cortico-cortical drives delivered to the modeled column. ii) Simulated spikes from cell units of the model during one example trial. iii) The average model dipole activity and averaged dipoles in each model layer (thin gray lines indicate single-trial dipole activity, thick lines indicate the all-trial average). iv) Depiction of which drive parameters in the model were optimized from a previous simulation.

*Current dipole simulation.* The net current dipole is simulated by summing the intracellular current flow in the pyramidal neuron dendrites directed parallel to the apical dendrite. A multiplicative scaling factor is applied to simulated net current dipole to estimate the number of cells contributing to the recorded signals, under the assumption that the recorded macroscale signals reflect nearly synchronous pyramidal neuron activity across larger populations of cells than those simulated. Here, we used a scaling factor of 150. Though the scaling factor relates roughly to the number of neurons producing the dipole, this correlation is indirect in the context of sensor-level data, as discussed further below. Additionally, raw evoked responses were smoothed by convolving with a 30ms Hamming window to account for spatiotemporal averaging that occurs when recording the response over a larger heterogeneous network in the data as compared with the model.

*Exogenous drive.* There are two types of exogenous excitatory synaptic drive that can activate the local network. One drive is designed to represent feedforward input from the lemniscal thalamus that targets Layer 4 (not explicitly modeled) and is then relayed to the inhibitory neurons and pyramidal neuron basal (or proximal) dendrite compartments in Layers 2/3 and Layer 5, referred to as Proximal drive in **Figure 1**. The other drive represents feedback input from either higher-order cortex or non-lemniscal thalamus and targets inhibitory neurons in Layer 2/3 and the distal dendrites of the Layer 2/3 and Layer 5 pyramidal neurons, referred to as Distal drive in **Figure 1**. In practice, users define patterns of pre-synaptic spike trains representing exogenous drive from external sources, and these spikes activate excitatory synapses in the local network. Stochasticity is built into the timing of the exogenous drive such that on each simulation “trial” the time of the input spike (or spikes) is chosen from a Gaussian distribution with a user-defined mean input time and standard deviation (SD). For each condition we simulated 50 trials, in alignment with the average number of trials yielded for each condition in the empirical data. Individual trial and averaged data are shown. The exogenous drive together with induced local network spiking generates current flow up and down the pyramidal neuron dendrites to produce the net current dipole signal. Although the exact sources of external drive remain abstract in the model, we discuss potential sources of these drives based upon structural and cytoarchitectural principles in the Discussion.

*Using HNN to estimate sensor-level data.* As described above, HNN is designed to simulate current dipole activity from a single source in the brain. Here, we applied ICA to the sensor-level signals as described above to estimate the activity from a single neural generator. This method does not provide information about the anatomical location of the source, and the units are in mV rather than current dipole units of nAm.

The primary assumptions in comparing HNN current dipole output to these ICA applied sensor signals is that that signal reflects activity from a single brain source, here assumed to be in mPFC. As such, positive and negative deflections in the sensor level signal can be related to the primary current generator coming from intracellular current flow up and/or down the pyramidal neuron dendrites (see also Sliva et al., 2018, where HNN was used to study sensor level data). The direction of the current at any point in time can thereby be inferred with source localization techniques when structural scans are available. However, since we do not have access such scans, here we make the assumption that the first positive peak (P2) reflects a feedforward input that drives current flow up the pyramidal neuron dendrites; this assumption is consistent with prior studies examining the sequence of spiking activity across the cortical layers (Sajad et al., 202; see further discussion in the Results section). For completeness, we also examined an alternative model where the first peak reflected a downward-directed current; this simulation did not provide a good fit to our data. Further, we examined the possibility that the FC-ERP might be generated by two different sources in the mPFC rather than a single source; these results were consistent with the main conclusions from our study. See the Results section for a detailed examination and discussion of each of these alternatives.

*Process for hypothesis testing and parameter estimation in HNN.* HNN contains a complex dynamical systems model representing a neocortical column under exogenous driving influences. Due to the large-scale nature of the model, which has hundreds of parameters, in practice users leave most of the parameters fixed and adjust only a user-defined subset of parameters based on hypotheses motivated by prior studies and literature. The graphical user interface and quantification of the root mean squared error (RMSE) between simulated and empirical evoked response waveforms allows users to examine if and how adjustments in these parameters can account for recorded current dipole source data.

Given the HNN neocortical template model (**Figure 1**, Extended Data Figure 3-2), all simulations begin by “activating” the network with some type of assumed exogenous drive that depends on the experimental conditions. For evoked responses, as studied here, it is hypothesized based on prior studies in sensory cortex that the local network receives a sequence of initial feedforward thalamocortical input (Proximal 1) followed by a cortico-cortical feedback input (Distal 1) and subsequent re-emergent thalamocortical feedforward input (Proximal 2; e.g., see Jones et al., 2007; Kohl et al., 2022; and schematic illustrations in **Figure 3Aiv, 3Biv**). These studies were used to establish the default input pattern distributed with HNN, which reproduces a tactile-evoked response. Here, we built upon these studies to propose similar mechanisms of generation for the FC-ERP, under assumptions that there is a consistent canonical structure and layer-specific organization of external inputs across cortical regions (see Discussion; e.g., Barbas, 2015; E. G. Jones, 1998; Vogt et al., 1987). Moreover, prior work has demonstrated that excitatory and inhibitory cells in Layers 2/3 and Layer 5 of primate prefrontal cortex spike across early and late windows following task-related events in the stop-signal task (Sajad et al., 2022), in alignment with this default sequence of inputs. Given this default sequence, we estimate the number, timing, and maximal conductance of these drives to best account for the recorded data. Initial and several alternative hypotheses on the parameters of these drives are tested to identify configurations that reproduce the best fit to data (i.e., smallest RMSE). The best fit model provides targeted predictions on the multiscale mechanisms creating the current dipole source signal that can guide further follow up testing with invasive recordings or other imaging modalities (e.g., see Bonaiuto et al., 2021; Sherman et al., 2016).

The process for parameter estimation begins by starting with the hypothesized sequence of drives, using the drive parameters distributed with the software, and manually hand-tuning the parameters defining the timing and strength of these drives to get an initial close representation to the current dipole waveform, or, small RMSE between the empirical current dipole and simulated trial-mean current dipole. Once an initial close representation to the data is found, automated parameter estimation algorithms distributed with the software that leverages the COBYLA algorithm (Powell, 1994) can be used to estimate parameters (within a defined range; here considered to be no more than 300% for synaptic weight changes to avoid biologically implausible changes) that produce the smallest RMSE between the simulated and recorded data (i.e., the “best fit” to the data). Several runs of algorithmic optimization were conducted when fitting the default HNN parameters to the successful Stop data and the successful Stop model parameters to failed Stop data. Following algorithmic fitting, we manually tested whether synaptic changes made during optimization were necessary to improve model fit and eliminated changes that did not improve the fit of the average dipole to waveform characteristics. The pre-tuned HNN model and the described process for hypothesis testing and parameter estimation has been successfully applied in several prior studies of sensory evoked responses (Jones et al., 2007, 2009; Kohl et al., 2022; Law et al., 2022; Sliva et al., 2018; Thorpe et al., 2021). This study is the first to apply this process to study evoked responses from frontal cortex.

### Horse-race model hypotheses and implications for neural signatures of the Stop process

As briefly discussed in the Introduction, theorized links between inhibitory processes and neural signatures such as the FC-ERP in the SST have been motivated in large part by assumptions and predictions of the horse-race model of SST behavior (Logan & Cowan, 1984a; Verbruggen & Logan, 2009). This model assumes that the outcome of a given Stop trial is determined by the winner of a race between an underlying Go and Stop process, which begin following their respective eliciting stimuli. Although the Go process begins earlier, it is slightly slower than the Stop process. The original iteration of this model (Logan & Cowan, 1984a) examined variations in the Go process speed (i.e., length or RT) assuming invariant Stop process speed, implying that whether or not a Stop was successful relied heavily on SSD. However, this is not reflected in neural dynamics, which indicate both the Stop and Go timing matters when determining the success of stopping (e.g., Schmidt et al., 2013). Critically, historic and more current iterations of the horse-race model assume that the success of stopping is not determined by the strength of the Stop process, but rather its timing relative to Go. Given the variance of the speed of the Go process, an earlier start for the Stop process is advantageous and more likely to result in a successful Stop. These predictions parallel the characteristics of the FC-ERP’s P3 deflection observed during successful and failed Stops: the amplitude of the P3 does not vary significantly by stopping accuracy, but onsets significantly earlier in successful versus failed Stops.

## Results

### P3 onset occurred earlier in successful compared to failed Stops in a Stop-Signal Task

Participants performed a SST in which Stop-Signals (red arrows) were presented following the Go signal on one–third of trials. Grand-average ERPs (**Figure 2**) were calculated at electrode FCz, time-locked to the Stop-Signal for Stop trials or to the current Stop-Signal delay in the staircase for Go trials. Consistent with prior studies (i.e., de Jong et al., 1990; Enriquez-Geppert et al., 2010; Kok et al., 2004; Wessel & Aron, 2015), the FC-ERP observed during Stop trials had several prominent waveform features: 1) the P2, an early positive-going deflection that occurred approximately 150ms after the Stop-Signal, 2) the N2, a negative-going deflection that occurred approximately 200ms after the Stop-Signal, and 3) the P3, a very large and sustained positive-going deflection that peaked at approximately 300ms following the Stop-Signal. A prominent P3 deflection was visible on Stop but not Go trials (**Figure 2C**). The P3’s *onset* occurred significantly earlier in successful compared to failed Stop trials (*t*(233) = –4.94, *p* < .0001, *d* = 0.29). The amplitude of the P3 did not differ between conditions (*t*(233) = 0.84, *p* = 0.40, *d* = 0.02). In addition, P2 peak amplitude was larger on failed Stop trials than successful Stop trials (*t*(233) = –8.53, *p* < .0001, *d* = 0.44), while the N2 peak amplitude was not significantly different between conditions (*t*(233) = –1.68, *p* = 0.09, *d* = 0.07). However, N2 peak latency was significantly earlier in successful Stops than in failed Stops (*t*(233) = –5.09, *p* < .0001, *d* = 0.34). The presence of the N2 and P3 peaks and their relation to the success of Stop trials here replicated prior work. Compared to the N2/P3 complex, the P2 is not typically theorized to relate to the activity of the Stop process itself per se, but represents a canonical feature of the FC-ERP that impacts preceding network dynamics and was therefore modeled alongside the N2 and P3 features in our HNN models (see next section).

### HNN predicted the Stop-Signal FC-ERP could be generated by a sequence of exogenous proximal and distal thalamocortical drives

A main goal of this study was to use HNN to simulate the thalamocortical and cortico-cortical mechanisms generating differences in the FC-ERP between successful and failed Stop trials, and specifically to evaluate whether predicted mechanisms of P3 onset latency differences are consistent with race model-derived assumptions of an earlier Stop process onset in successful Stops. To begin, we modeled the mechanisms by which the FC-ERP could be generated, focusing first on recapitulating the key features of the Stop-Signal-locked FC-ERP observed during successful Stop trials: namely, the polarity and timing of the P2, N2, and P3 peaks. For more logic behind hypothesis testing with HNN and a discussion of how this was accomplished procedurally, we refer the reader to the Methods.

We began with the HNN default evoked response parameters distributed with the software that define a sequence of thalamocortical and corticol-cortical feedforward and feedback input to the local network through Proximal and Distal projection pathways (e.g., **Figure 3Aiv**). We first hand-tuned the mean onset latency and scaling factor (see Methods) of these inputs to produce an initial fit to the FC-ERP features. Additionally, while the HNN default model assumes one incoming spike on each trial for the proximal drive, we found that two incoming spikes were needed to reproduce the long, sustained P3 deflection of the FC-ERP (see **Figure 3-2 in the Extended Data**). Once initial agreement to the ERP shape was found via hand tuning, we conducted algorithmic optimization of a subset of parameters in order to minimize the RMSE between the model output and recorded data. The parameters defining the exogenous thalamocortical and cortical-cortical drives were targeted for optimization and included: 1) the timing and standard deviation across trials of the driving spikes defining each input to the model and 2) the synaptic weights (i.e., maximal conductance) of AMPA and NMDA receptors on the target pyramidal and basket cells for each input, while all other parameters in the model (i.e., local network parameters, etc.) remained set to default values. The adjusted parameters and the degree to which they were changed from the HNN ERP default parameters to produce the FC-ERP in successful Stop trials (as measured by a small RMSE, e.g., **Figure 3Aiii**) are reported in **Extended Data** Figure 3-3. Of note, remarkably few changes needed to be made to HNN default parameters in order to fit the FC-ERP, suggesting ERPs from the frontal cortex are constrained by similar canonical anatomical and physiological features as those in sensory cortex (including somatosensory and auditory cortex; Jones et al., 2007; Kohl et al., 2022).

The optimized HNN model predicted that an initial thalamocortical (Proximal 1) drive at 115ms, a cortico-cortical (Distal 1) drive at 190ms, and a re-emergent thalamocortical (Proximal 2) drive at 306ms led to the FC-ERP observed following Stop-Signals during successful Stop trials (RMSE = 0.41; **Figure 3A**). An alternate initial model with a distal-proximal-distal drive sequence produced deflections that were too peaked to fit the positive-going features of empirical FC-ERPs even after optimization of model parameters (see **Extended Data Figure 3-4**).

To establish an understanding of how the described sequence of drive creates the final successful Stop FC-ERP (**Figure 3A**), we here review a few key model concepts detailed in prior studies (e.g., see Neymotin et al., 2020; see also Methods). Synaptic excitation via proximal and/or distal drive pushes current away from the location of the synapse, while inhibition pulls current toward the location of the synapse. Backpropagation of action potentials creates upward current flow and dendritic calcium spikes generate downward current flow (e.g., see Law et al., 2022). As such, in the simulation shown, the initial proximal drive combined with backpropagation of pyramidal neuron spiking activity created upward current flow and the positive P2 peak, while the subsequent distal drive and strong synchronous somatic inhibition created downward current flow and the negative peak (N2). The second proximal drive generated a subsequent bout of upward current to create the positive P3 peak.

A spiking histogram of each cell in the network is shown for an example trial in **Figure 3Aii**. The inlaid box in the average dipole plot (**Figure 3Aiii**) displays the contribution to the net current dipole from each Layer of the model (see also extended data **Figure 3-5** for non-smoothed results). Layer 5 contributed more to the averaged dipole because dendrites of pyramidal cells in this layer of the model were longer, resulting in a greater contribution to the net current flow.

### HNN predicted that proximal drive associated with P3 onset occurred earlier in successful Stops compared to failed Stops, in support of race model predictions

We next tested the race model-derived hypothesis that the difference in P3 onset during successful Stops could be accounted for by Proximal 2 drive that arrives earlier (i.e., earlier mean onset time of drive to cortex) but is not effectively stronger (i.e., increases in excitatory synaptic weights) than in failed Stops. The process for fitting the HNN model to the failed Stop FC-ERP was to start with the model fit to the *successful* Stop FC-ERP and conduct algorithmic optimization of a targeted subset of parameters to produce a close representation to the failed Stop FC-ERP (see Methods). Once more, we optimized parameters associated with the timing, variance, and synaptic weights of exogenous proximal and distal drives (see previous section). However, for the Proximal 2 drive, only the timing and variance of the inputs were optimized, with synaptic weights held constant to specifically test the hypothesis that the effective strength of this input does not need to change to account for differences in P3 latency.

Consistent with race-model based predictions of a *later* but not *weaker* latent Stop process in failed Stops, in the optimized failed Stop model, the onset of the Proximal 2 drive was later and the strength of the drive did not need to be adjusted. Specifically, the best fit model (RMSE = 0.44; **Figure 3B**) predicted a mean onset time of 325ms for Proximal 2 drive, approximately 19ms later than in successful Stops (see **Table 1**).

**Table 1.**
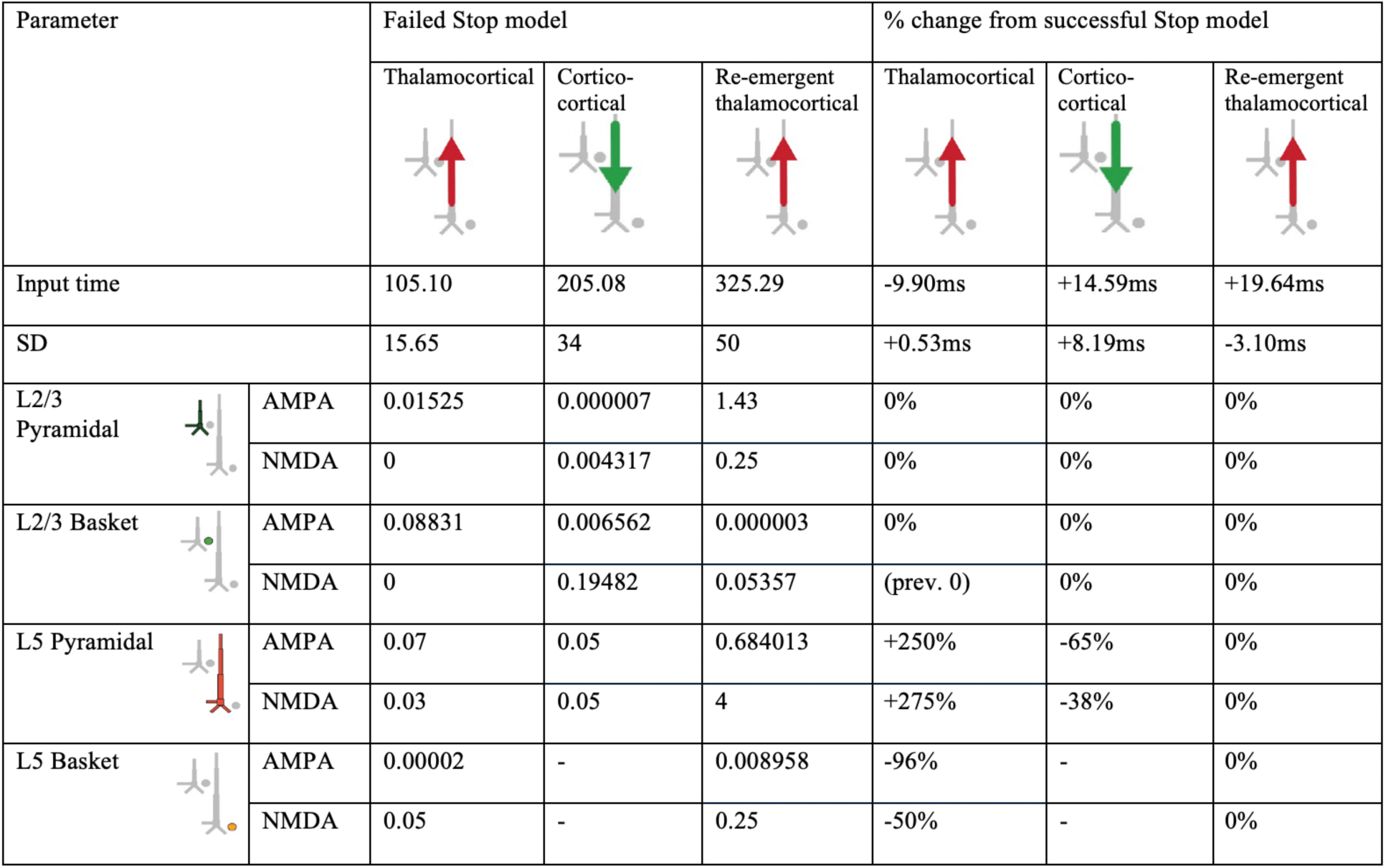
HNN model parameters for the failed Stop FC-ERP model and percentage changes from the successful Stop model parameters after optimization.

| Parameter |  | Failed Stop model |  |  | % change from successful Stop model |  |  |
| --- | --- | --- | --- | --- | --- | --- | --- |
|  |  | Thalamocortical | Cortico-cortical | Re-emergent thalamocortical | Thalamocortical | Cortico-cortical | Re-emergent thalamocortical |
| Input time |  | 105.10 | 205.08 | 325.29 | -9.90ms | +14.59ms | +19.64ms |
| SD |  | 15.65 | 34 | 50 | +0.53ms | +8.19ms | -3.10ms |
| L2/3 Pyramidal | AMPA | 0.01525 | 0.000007 | 1.43 | 0% | 0% | 0% |
|  | NMDA | 0 | 0.004317 | 0.25 | 0% | 0% | 0% |
| L2/3 Basket | AMPA | 0.08831 | 0.006562 | 0.000003 | 0% | 0% | 0% |
|  | NMDA | 0 | 0.19482 | 0.05357 | (prev. 0) | 0% | 0% |
| L5 Pyramidal | AMPA | 0.07 | 0.05 | 0.684013 | +250% | -65% | 0% |
|  | NMDA | 0.03 | 0.05 | 4 | +275% | -38% | 0% |
| L5 Basket | AMPA | 0.00002 | - | 0.008958 | -96% | - | 0% |
|  | NMDA | 0.05 | - | 0.25 | -50% | - | 0% |

While the timing of the Proximal 2 drive was the target of our hypothesis testing with this model, we note that synaptic weight changes occurred in the earlier Proximal 1 and Distal drives to account for differences across conditions in the earlier P2 and N2 peaks. The larger P2 amplitude in failed Stops emerged from an earlier and stronger (Layer 5 pyramidal weights increased, basket weights decreased) Proximal 1 drive. Although no significant differences in N2 amplitude were observed on average in our data, the failed Stop model predicted both a later and weaker (Layer 5 AMPA and NMDA weight reductions) Distal drive during failed Stop trials. These changes were necessary to compensate for differences in the earlier Proximal 1 drive underlying the P2 deflection, which in our simulations generated a different network state at the time of the N2 across conditions. Of note, the patterns of layer-specific spiking activity were similar across the successful Stop and failed Stop models, with the notable exception of more bursty spiking activity early in the trial in the Layer 5 pyramidal neurons in the failed Stop model (**Figure 3Biii**). One interpretation of this early increase in spiking is that it may increase output from frontal cortex to subcortical and motor areas, ultimately erroneously facilitating movement in failed Stop trials.

### Alternative models of P3 onset mechanisms could not simulate differences in P3 latency across Stop accuracy conditions

To avoid model bias stemming from our main hypothesis about the cause of the P3 onset latency difference across conditions, we also simulated two alternative hypotheses of mechanisms that could potentially account for the later P3 onset in failed Stops. Both alternatives tested whether decreases in the effective *strength* of re-emergent thalamocortical (Proximal 2) inputs could account for the later P3 onset, as opposed to a later onset time.

In one alternative, we decreased the effective strength of the Proximal 2 input by reducing the number of spikes representing this drive from two to one to simulate a weaker underlying Stop process in failed Stop trials. We then optimized the conductance strength (i.e., synaptic weights) and timing parameters of all input drives – except for the timing of Proximal 2 – to fit the recorded ERP (see Methods). The best fit waveform simulated with this alternative mechanism produced a worse fit to the Stop-Signal locked ERP than before, such that the amplitude of the P3 was too small and the onset of the P3 too early in failed Stops (see boxes in **Figure 4Aiii, compare to Figure 3Biii**). This was due to decreased proximal drive and an overall decrease in L5 pyramidal neuron firing (RMSE = 0.77; compare spike histograms in **Figure 4Aii** and **Figure 3Bii**).

**Figure 4.**
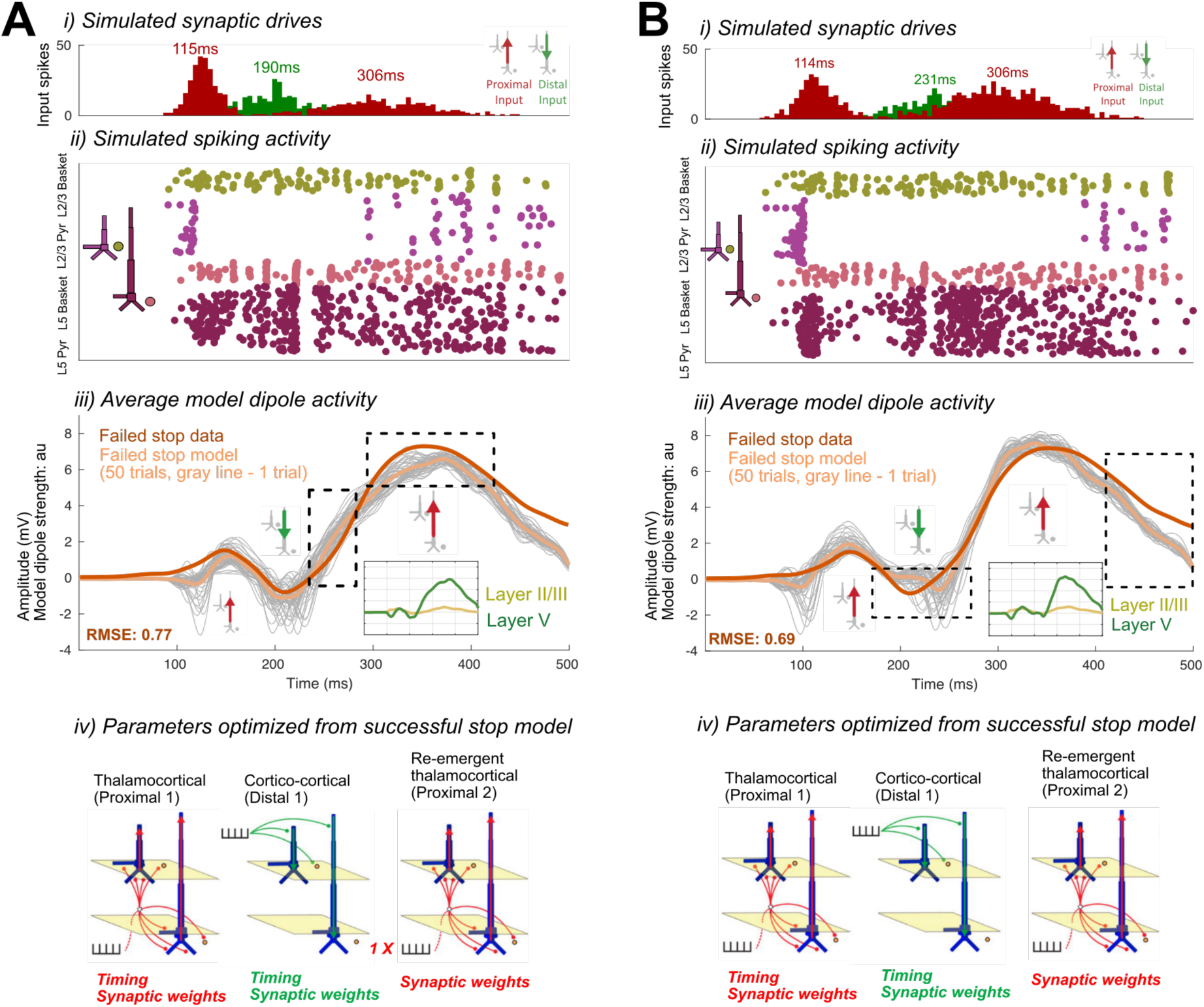
Alternate HNN models of re-emergent thalamocortical (Proximal 2) drive condition differences in the failed Stop FC-ERP. A) shows the alternate model in which the re-emergent thalamocortical (Proximal 2) drive contains one spike instead of two. B) shows the alternate model in which the weights and timing of all drives were allowed to change, except for the timing of the re-emergent thalamocortical drive (Proximal 2), which remained fixed during optimization. i) All-trials histograms of thalamocortical and cortico-cortical drives delivered to the modeled column. ii) Simulated spikes from cell units of the model during one example trial. iii) The average model dipole activity and averaged dipoles in each model layer (thin gray lines indicate single-trial dipole activity, thick lines indicate the all-trial average). iv) Depiction of which drive parameters in that model were optimized from a previous simulation.

**Figure 5.**
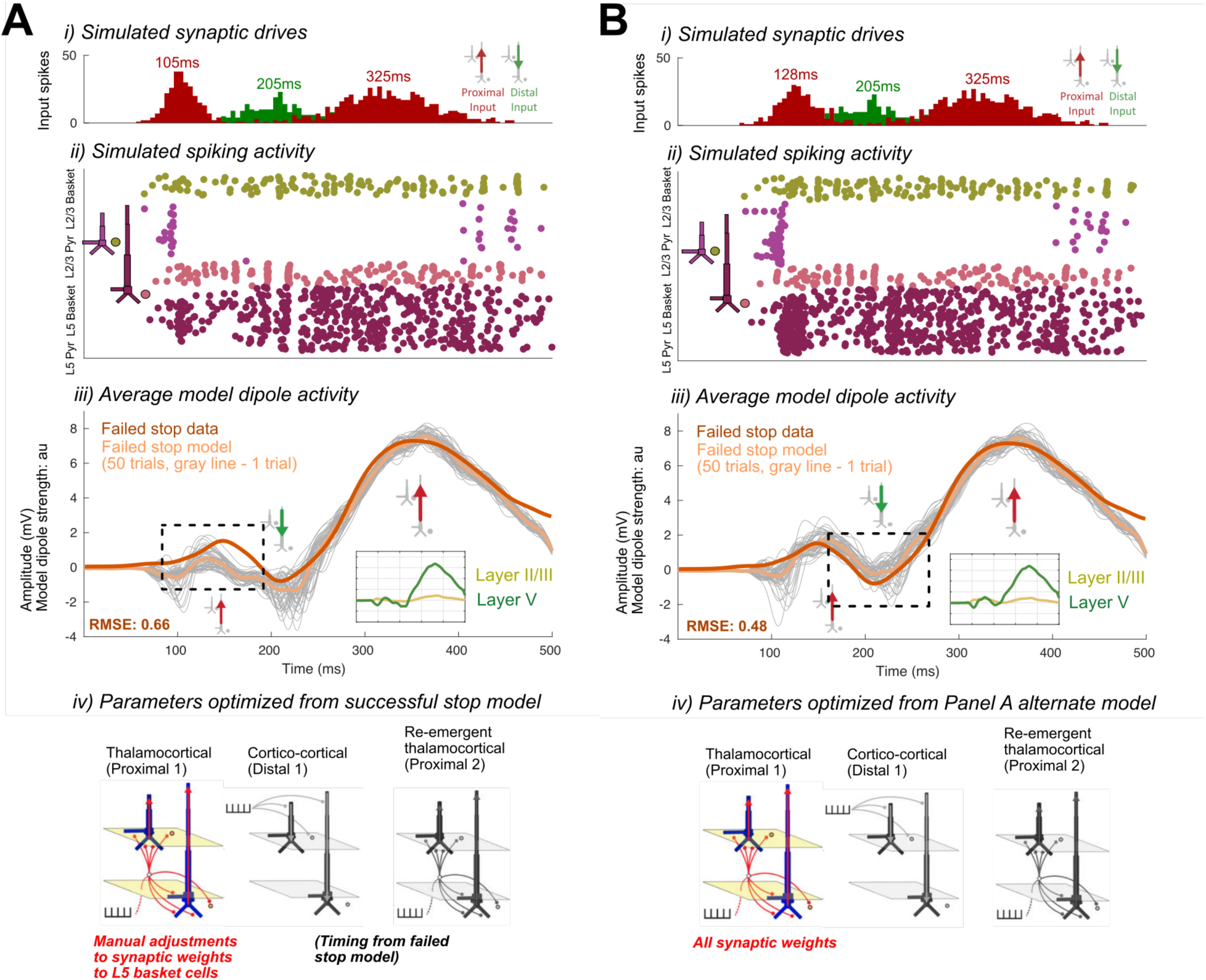
Alternate HNN models of cortico-cortical (Distal 1) drive condition differences in the failed Stop FC-ERP. A) shows the alternate model in which all model parameters were reverted to the successful Stop model parameters and Layer 5 basket cell synaptic weights were manually increased to simulate somatic inhibition. B) shows the alternate model in which optimization of the timing and all of the synaptic weight parameters of the thalamocortical drive (Proximal 1) was conducted, starting from the parameters of the model in A. i) All-trials histograms of thalamocortical and cortico-cortical drives delivered to the modeled column. ii) Simulated spikes from cell units of the model during one example trial. iii) The average model dipole activity and averaged dipoles in each model layer (thin gray lines indicate single-trial dipole activity, thick lines indicate the all-trial average). iv) Depiction of which drive parameters in that model were optimized from a previous simulation.

In the second alternative, rather than assuming the effective strength of the Proximal 2 input was weaker in the failed Stop model, we tested if a change in the conductance strength of this drive (in either direction) instead of its timing could also fit the data. More specifically, as in the first alternative, we optimized the conductance strength (i.e., synaptic weights) and timing parameters of all input drives except for the timing of Proximal 2, albeit in this case without reducing the number of drive spikes. Though the best fit model recapitulated some of the features of the empirical waveform (especially waveform components preceding P3), it could not fit all of the waveform characteristics as well as before (compare features highlighted with boxes in **Figures 4Biii and 3Biii**). First, the N2 deflection was later and larger in magnitude than the data, due to a later shift in the Distal drive, which did not overlap with the preceding thalamocortical Proximal 1 drive. This feature was present on nearly all trials (see gray lines in **Figure 4Biii** for individual trial simulations) as opposed to only some trials in the previous timing-change model (**Figure 3Biii**). Second, the late tail of the ERP (>400ms) returned too quickly to baseline, due to a reduction in Layer 5 pyramidal neuron spiking (RMSE = 0.69; compare spike histograms in **Figure 3Bii** and **Figure 4Bii**).

### Alternative models of P2 and N2 dynamics

We so far have focused on the simulated mechanisms predicted to underlie the difference in the timing of P3 onset in failed and successful Stop trials. However, there are also differences in earlier FC-ERP peaks, such that in failed Stops the P2 deflection is larger and the N2 deflection appears smaller (although this peak amplitude difference is not statistically significant in our data sample; **Figure 2C**). Our best-fitting model (see **Figure 3B**) also makes specific predictions about the changes to incoming drives that simulate these earlier differences. Specifically, the synaptic weights of the thalamocortical (Proximal 1) and cortico-cortical (Distal 1) drive producing the P2 and N2 became stronger and weaker in the failed Stop model, respectively. The weaker Distal drive compensated for a stronger Proximal 1 drive to maintain a nearly constant, albeit slightly smaller, N2 magnitude.

Given that the model predictions about the generation of these early P2 and N2 changes were a consequence of automated optimization, rather than a systematic exploration of specific hypotheses, we examined possible alternative mechanisms of P2 and N2 that could involve changes to Proximal 1 drive (generating P2) while requiring minimal changes to Distal drive (generating N2). Within HNN’s framework, a reduced efficacy in Distal drive and consequently smaller-amplitude N2 could arise from two primary mechanisms: 1) a reduction in distal depolarizing currents, such as due to increased synaptic weights of the Distal drive onto pyramidal cells, or 2) hyperpolarization of the pyramidal cell soma, resulting from inhibition produced in response to a preceding Proximal drive. Both of these mechanisms reduce downward current flow in apical dendrites during Distal drive. Our timing-change model of failed Stops (**Figure 3B**) predicts that the smaller N2 results from the former mechanism. Here, we test whether the latter could produce this same condition difference as well.

First, we tested if the compensatory reduced efficacy of Distal drive in the failed Stop model could also result from increased pyramidal inhibition at the time of the N2 peak. An increase in inhibition could emerge from changes in the preceding thalamocortical (Proximal 1) dynamics (e.g., Kurotani et al., 2008; Pouille & Scanziani, 2001), and specifically from an increase in the efficacy of the Proximal 1 drive onto the inhibitory cells that would in turn increase somatic inhibition of pyramidal cells. To test this, we reverted all the synaptic weights for the Proximal 1 and Distal drives back to the successful Stop model weights and then manually increased the synaptic conductance weights of thalamocortical (Proximal 1) drive to L5 Basket cells by rounding up the Proximal 1-Layer 5 basket AMPA weight parameter (from 0.000561 to 0.001, 79% increase) and doubling the Proximal 1-Layer 5 basket NMDA weight parameter (0.1 to 0.2, 100% increase). This led to a flatter and smaller amplitude P2 deflection that was not consistent with the empirical FC-ERP (RMSE = 0.66; compare features highlighted with box in **Figure 5Aiii** to similar features in **Figure 3Biii**).

We then expanded this exploration by starting from the increased Proximal 1 weights to the basket cells in **Figure 5A** and ran automated optimization on the timing, variation, and all of the synaptic weight parameters of the Proximal 1 driv*e.* The resulting model simulated a better fit to the empirical FC-ERP (**Figure 5B**), even though it did not fit the data quite as well as our original failed Stop timing-change model (RMSE = 0.48 versus 0.44 in **Figure 3B**), since the timing and amplitude of the N2 peak were slightly misaligned. Compared to the best-fitting failed Stop model in **Figure 3B**, in **Figure 5B** the mean timing of the Proximal 1 drive is slightly later, the synaptic conductances for Proximal 1 onto the inhibitory cells is stronger while the synaptic conductance onto the pyramidal cells is the same, and the synaptic conductances for Distal drive to the excitatory is stronger because they remained set at the successful Stop model strengths (compare **Table 1** with **Extended Data** Figure 5-2). Such a small difference in the fit of the two models in **Figure 3B and 5B** suggests that the simulated mechanism in either model could underlie empirical condition differences in the early FC-ERP waveform components (P2/N2). A key distinction in Proximal 1 dynamics from the successful Stop model (**Figure 3A**) in both failed Stop models (**Figure 3B** and **5B**) is the fact that this drive is stronger, generating more Layer 5 (**in Fig. 3Bii** and **Figure 5Bii**) and Layer 2/3 pyramidal cell spiking (**in Figure 5Bii**), which would lead to differences in the strength of early signaling to downstream regions. Model parameters for all tested alternative models are listed in **Extended Data Figures 4-1, 4-2, 5-1, and 5-2**.

### Modeled mechanisms are conserved when simulated with two sources

As discussed in the Methods section, HNN is designed to simulate current dipole activity from a single source within the brain. However, despite efforts to functionally localize the FC-ERP to mPFC generators (see Methods) this assumption cannot be met entirely when modeling sensor-level EEG activity. Therefore, we also examined the possibility that the FC-ERP might be generated by two different sources in the mPFC, whose activity sum together, rather than a single source. We also examined if the predicted mechanistic difference in successful vs failed Stop trials still holds with the two-source model (see Extended Data Figure 3-6).

To accomplish this, we first artificially split the empirical FC-ERP between the N2 and P3 deflection, assuming the earlier and later parts of the waveform come from two different underlying sources. Then, beginning with the parameters used for the successful Stop FC-ERP, we ran optimization on the parameters accounting for the P2/N2 (i.e., Proximal 1 and Distal drive) and P3 (i.e., Proximal 2 drive) deflections separately, fitting two different models (neocortical sources) to the two parts of the waveform. This yielded two models that accurately reproduced the (1) P2 and N2 deflections and (2) the P3 deflection, respectively. Further, when summed together the output from the two models reproduced the entire time course of the successful Stop FC-ERP, suggesting that it is not necessary that these individual waveforms all arise from the same underlying source (see Extended Data Figure 3-6**, A-C**).

To examine if the predicted mechanisms underlying successful vs failed Stops still hold, we next optimized these two models to the failed Stop FC-ERP, where the empirical data was similarly artificially split into early and late components. To be consistent with our prior hypotheses and simulations, we allowed all parameters of the first column’s model – which included the Proximal 1 and Distal drives – to vary during optimization, but only allowed the timing parameters in the second model – which contained the Proximal 2 drive – to vary. As before, we found that changes to timing and strength (in the same direction) of the parameters associated with the first Proximal and Distal drive were needed to account for early differences in the P2 and N2, but that a timing change alone in the second Proximal drive could account for the later P3 in failed Stops. Again, when summed together the output from the two models reproduced the entire time course of the failed Stop FC-ERP (see **Extended Data Figure 3-6, D-F**). These results suggest the predicted mechanisms of successful vs failed stopping can hold even if the FC-ERP is generated by two sources.

### Summary of HNN model-predicted differences between mechanisms of the successful and failed Stop FC-ERP and implications for behavior

In this section so far we have described the logic we followed to optimize the successful and failed Stop FC-ERP models based on *a priori* hypotheses and *post-hoc* investigation of simulated alternative mechanisms. Our investigation resulted in a number of specific model-based predictions on the mechanisms of FC-ERP generation in each condition, as shown in **Figure 3A** and **Extended Data** Figure 3-2 for successful Stops and **Figures 3B/5B**, **Table 1**, and **Extended Data** Figure 5-2 for failed Stops. Here, we summarize these predictions (**Figure 6)** and their implications for SST behavior.

**Figure 6.**
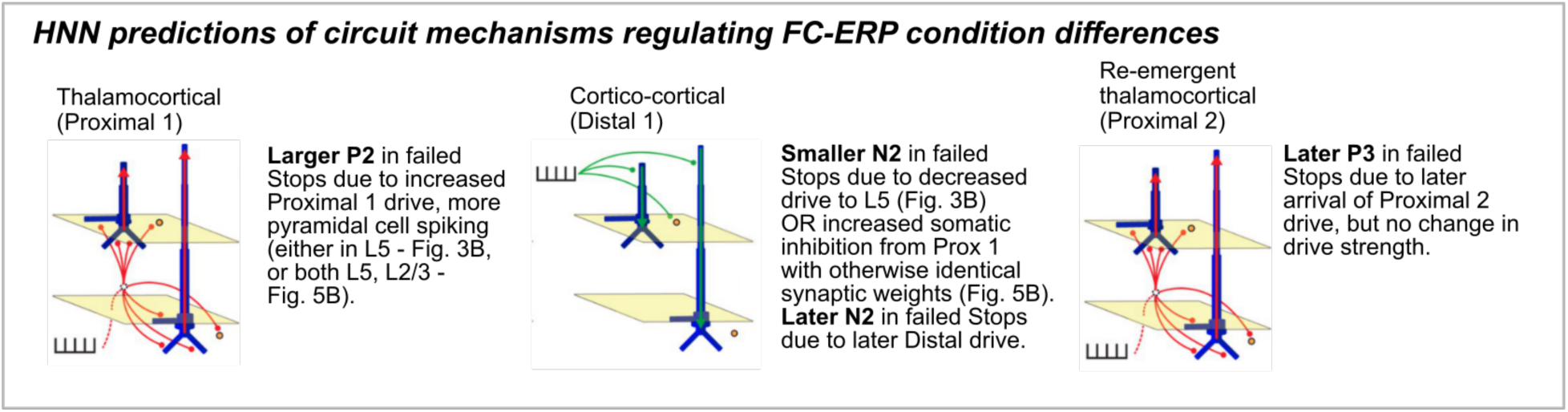
Diagram of HNN-predicted mechanistic differences between successful and failed Stop FC-ERPs.

*P2 deflection*. Modeled thalamocortical (Proximal 1) drive to the pyramidal neurons was stronger in failed compared to successful Stops, driving more early spiking at the time of the P2. Proximal 1 drive was stronger to Layer 5 pyramidal cells in both failed Stop models shown in **Figure 3B** and **Figure 5B**, and in the case of **Figure 5B** also to the Layer 2/3 pyramidal cells, producing a larger positive current deflection in the model dipole which accounted for a higher-amplitude P2 in failed Stop trials. A prediction of increased firing in mPFC neurons early following the Stop-Signal in failed Stops implies that regulation of this early activity (which could be related to a prepotent Go response) is critical for stopping success.

*N2 deflection*. Modeled cortico-cortical (Distal) drive producing the N2 in failed Stops occurred later than in successful Stops in both models. In the failed Stop model in **Figure 3B**, the Distal drive compensated for a larger P2 through a decrease in drive to Layer 5 pyramidal cells, which decreased spiking. In the failed Stop model in **Figure 5B**, lingering somatic inhibition of Layer 5 pyramidal cells from Proximal 1 drive counteracted the same synaptic Distal drive weights as in successful Stops to produce a smaller N2 deflection and less spiking without a change in synaptic weights. These models predict that an earlier distal drive supports stopping success, and that successful Stops may also be facilitated by increased L5 spiking in the N2 time frame (**Figure 3B**; or, alternatively, that there is no difference in the strength of Distal drive between successful and failed Stops; **Figure 5B**).

*P3 deflection*. In both models of failed Stop trials, modeled re-emergent thalamocortical (Proximal 2) drive occurred on average later, but did not differ in effective strength, compared to successful Stop trials. This result implies that successful stopping relies upon tightly timed firing of Layer 2/3 and Layer 5 pyramidal cells, which are recruited earlier in successful Stops but do not otherwise differ in their activity compared to failed Stops.

## Discussion

We used HNN’s biophysical model of the canonical neocortical column under exogenous drive to model the cell and circuit mechanisms underpinning the Stop-Signal-locked FC-ERP. Motivated by prior studies on the mechanisms generating sensory evoked responses (Jones et al., 2007, 2009; Kohl et al., 2022), we demonstrated that a similar sequence of simulated external thalamocortical and cortico-cortical drives can produce the FC-ERP waveform during successful Stops. We then examined the mechanisms underlying FC-ERP differences in successful versus failed Stop trials to draw neural mechanistic parallels to the predictions of the behavioral horse-race model of the SST, which implies that an earlier onset of an otherwise equivalent Stop process would lead to a higher probability of stopping success. We specifically tested the hypothesis that re-emergent thalamocortical drives generating the P3 (whose onset is proposed to index the Stop process, see Introduction) arrive earlier in successful Stop trials, but are not effectively stronger, than in failed Stops. By also modeling several alternative hypotheses, our results ultimately supported and provided a detailed circuit-level mechanistic explanation for this prediction, as well as a novel interpretive framework for differences in the earlier P2/N2 deflections that may contribute to the Stop process.

### Novel predictions of drive sequence producing the FC-ERP are consistent with prior studies and current theories of motor inhibition

Our modeling results predicted that the FC-ERP during successful stopping was generated by a sequence of thalamocortical (Proximal 1 at ∼115ms), cortico-cortical (Distal 1 at ∼190ms), and re-emergent thalamocortical (Proximal 2 at ∼306ms) drives, which sequentially interacted with local network dynamics to produce the P2, N2, and P3 peaks. In the model, the origins of these drives to the neocortex are abstract, and assumed to originate in thalamus and higher order cortex (or non-lemniscal thalamus), representing “feedforward” and “feedback” drive (Neymotin et al., 2020). This model-derived sequence is similar to that shown for evoked responses in primary sensory areas (e.g., Jones et al., 2007; Kohl et al., 2022), albeit at different latencies.

Several of our model-based predictions about mechanistic underpinnings of the Stop-Signal-locked FC-ERP align with emerging theories of motor inhibition as a two-stage process (Diesburg & Wessel, 2021; Schmidt & Berke, 2017). The Pause-then-Cancel (PTC) model of motor inhibition assumes that successful action-stopping requires both a rapid, nonselective Pause and a slower, selective Cancel process. Increased neuronal activity associated with the erroneous responses made in failed Stops in the early post-Stop-Signal (i.e., P2) timeframe aligns with the suggestion that the successful downregulation of action output by Pause benefits the overall success of stopping in successful Stops. On the other hand, the P3 exhibits several characteristics in line with a possible signature of the Cancel process. Prior work has shown that P3 amplitude is larger when associated with the explicit instruction to stop an action as opposed to ignoring the stimulus (Dutra et al., 2018; Tatz et al., 2021; Waller et al., 2021). Our models predict the Stop-Signal P3 is generated by thalamocortical inputs that arrive later but are not effectively stronger in failed versus successful Stops, which would be in line with a Cancel process that is deployed (albeit unsuccessfully in failed Stops) in a top-down fashion in both contexts due to task instructions to stop after the Stop Signal.

The precise sources for the presumed thalamocortical and cortical-cortical drives to the frontal cortex generating the human FC-ERP are unknown. Based on cytoarchitectural principles (e.g., Barbas, 2015; Barbas et al., 2013; Xiao et al., 2009; Zikopoulos & Barbas, 2007) and tract-tracing research in non-human animal models, we here propose several potential thalamic and cortical sources for these drives based on known patterns of laminar innervation. This is intended to guide further work testing the underpinnings of the FC-ERP by providing an illustration of how these factors may guide hypothesis formation following HNN investigations. As an example, we focus on data from the mPFC, as previous work suggests that this area may contribute to the FC-ERP following the Stop-Signal in humans (e.g., Huster et al., 2011). The citations within this section are drawn from investigations of dysgranular midcingulate cortex, but we expect laminar innervation in other regions of mPFC to be conserved, albeit from different source locations. Possible afferent sources of the initial Proximal drive to dysgranular layers (producing the P2) are thalamic core cells in the ventral anterior (VA) nucleus of the thalamus (known to provide ascending motor information; Vogt et al., 1987), the parietal association cortices (Fillinger et al., 2017; Vogt & Pandya, 1987), or even visual areas such as V2 or V1 (Fillinger et al., 2017). The Distal drive to supragranular layers (producing the N2) may arise from non-lemniscal thalamic matrix cells originating in the anterodorsal (AM) or mediodorsal (MD) thalamic nuclei (Vogt et al., 1987; whereby sensory and limnic information is integrated, e.g., Barbas et al., 2011), directly from limbic regions such as the amygdala (as Barbas et al., 2011 demonstrated for OFC; potentially carrying information about reward/salience), or from higher-order cortices like the agranular insular cortex (AI; Vogt & Pandya, 1987; see **Figure 7**).

**Figure 7.**
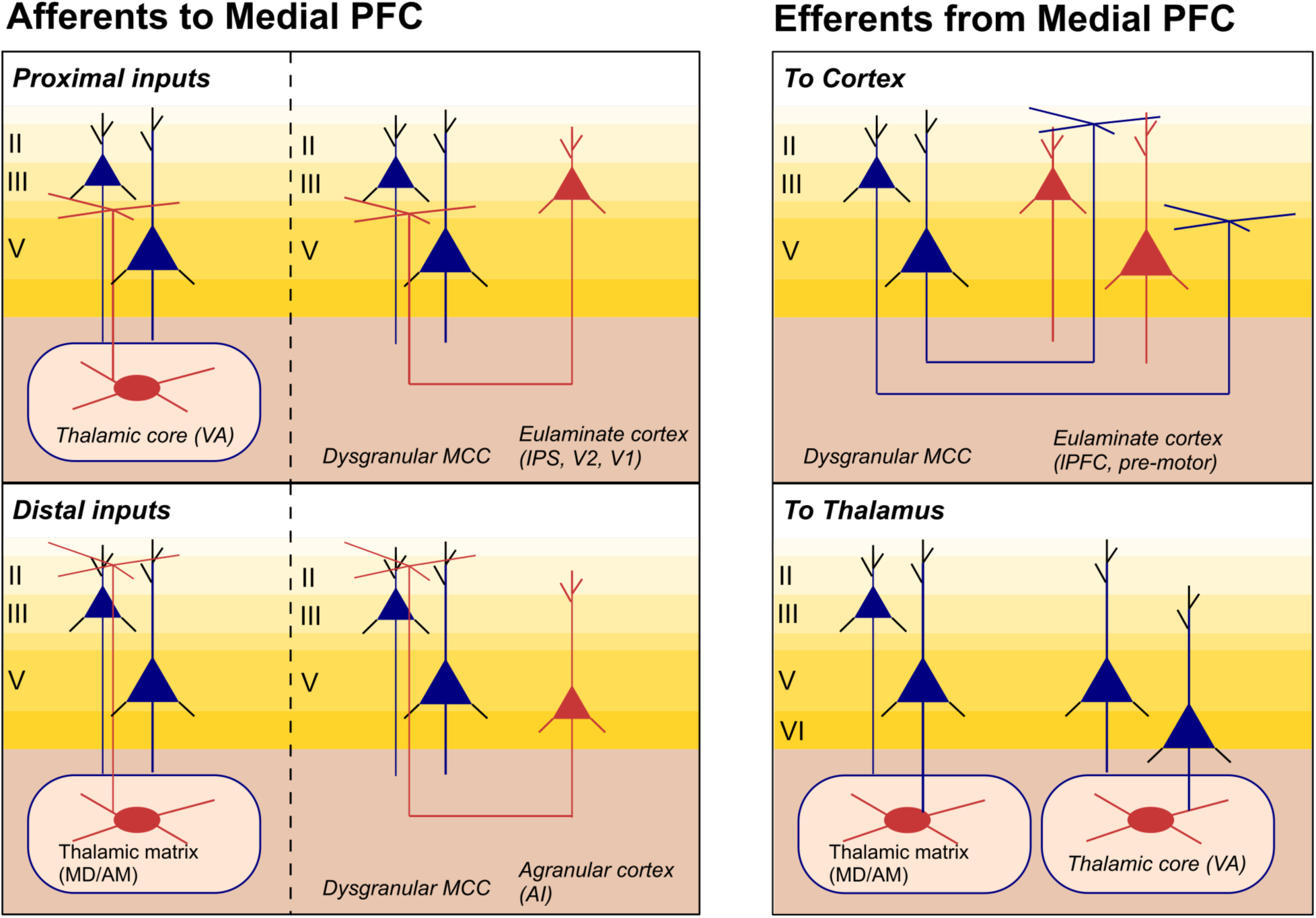
Proposed connections to and from mPFC. Proximal and distal connections to mPFC generators of the FC-ERP could come from thalamic core cells (in VA) and thalamic matrix cells (likely from MD or AM), or from cortical sources. Ascending sensory information constituted by proximal drive could come from eulaminate cortices such as the parietal association regions or even from the visual pathway. Feedback cortico-cortical information is more likely to come from even higher-order cortices such as AI. The FC-ERP could influence motor behavior by way of Layer 2/3 and 5 cortico-cortical to lower-order eulaminate cortices like the lateral pre-motor and pre-supplementary motor areas or via Layer 5 or Layer 6 signaling to the thalamic matrix or core, respectively. (Note that Layer 6 is not explicitly modeled in HNN.)

Efferent projections from mPFC to thalamic and/or cortical targets are also important to consider, as these projections provide possibilities for re-emergent thalamocortical signaling (which we assume produces the second proximal drive generating the P3 in our models) and ultimately contribute to network activity supporting the Stop process during successful action-stopping. Projections from Layer 2/3 and 5 of mPFC may target dorsal motor areas such as the premotor or pre-supplementary motor areas (Medalla et al., 2022), and Layer 5 and Layer 6 (which is not explicitly modeled in HNN currently) neurons are known to project back to both higher– (i.e., MD/AM) and lower-order (i.e., VA) thalamic nuclei, respectively (Domesick, 1969; Zikopoulos & Barbas, 2007), which could contribute to re-emergent thalamocortical signaling (see **Figure 7**).

### Model predictions help address debates on the role of N2 and P3 mechanisms in motor inhibition

An ongoing debate about relationships of FC-ERP features to underlying inhibitory processes in the SST is whether the N2 or P3 relates more directly to motor inhibition (see Wessel & Aron, 2015 and Huster et al., 2020). It has long been proposed that the P3 relates to inhibitory processes for the reasons discussed in the Introduction. However, the timing of the N2 peak also correlates with SSRT and occurs earlier in successful Stops (Anguera & Gazzaley, 2012; Huster et al., 2020; Senderecka, 2016; Senderecka et al., 2012). Our modeling reveals that it is likely the sequential and interacting dynamics of *several* circuit mechanisms generating Stop-Signal-locked ERP features that together support inhibitory control processes in the SST. Our models predicted that the N2 was generated by Distal input to superficial layers that generated downward currents followed by spiking in L5 pyramidal cells. In failed Stop trials, this drive occurred later and in one model (see **Figure 3B** versus **Figure 5B**) was effectively weaker than in successful Stop trials. This was because the N2’s timing and amplitude was impacted by the network state induced by initial thalamocortical (proximal) feedforward inputs, with lingering excitatory and inhibitory currents in both layers after the first proximal drive impacting the strength of the Distal drive. Hence, the dynamics regulating the first proximal drive not only play a crucial role in supporting action-stopping but continue to impact the network state when the drive producing the N2 arrives. This is also directly in line with work (discussed in the Method section) demonstrating the strong augmentation in mPFC that can arise from spike-timing dependent plasticity.

Setting aside the differences in strength in Distal drive between the failed Stop models, a key prediction of both is the later arrival of Distal input in failed Stops. In failed Stop trials, the re-emergent thalamocortical generating the P3 drive was also later than in successful Stops and may have failed to successfully support motor inhibition by arriving too late. This implies, given that both the N2 and P3 arrive earlier in successful Stops, that the precise signaling patterns from the mPFC to output regions in support of successful stopping may rely on the distal and proximal drives producing these deflections (and associated spiking) being appropriately timed and spaced with respect to each other.

In summary, both the N2 and the P3 are generated by mechanisms that vary with the success of stopping and have the potential to impact stopping accuracy through communication with downstream regions. These results also highlight that the N2 and P3 are underpinned by inputs from different origins to different cortical layers. More work is needed to disentangle the separate but parallel contributions of these mechanisms and resultant network states to stopping. Because the precise processes indexed by the N2 and P3 are still subject to debate, these models and subsequent work may help the field generate new ideas about the computations each feature reflects.

### Relevance to error-related signals in inhibitory control tasks

Here, we focused on the Stop-Signal-locked FC-ERP, whose defining characteristics include the negative-positive N2/P3 complex. Several other inhibitory task contexts are known to elicit similar frontocentral negative-positive complexes including action errors (ERN/P3; Gehring et al., 1993), unexpected action outcomes (N2/P3; Iwanaga & Nittono, 2010), and unexpected perceptual events (N2b/P3; Courchesne et al., 1975). Though the multiscale mechanisms underlying these FC-ERPs is unknown, ICA-based EEG analyses suggest that some of these signatures may come from the same underlying functional sources (Dutra et al., 2018; Wessel et al., 2012; see also Wessel & Aron, 2017). Recent efforts have similarly used exogenous spike inputs to basal and apical dendrites of isolated excitatory cells to simulate mechanisms underlying the primate intracranial ERN in supplementary eye fields during a saccade countermanding task, with a specific focus on understanding the relationship to theta power (Herrera et al., 2023). Modeling results in that study were not able to fully articulate the relationship to scalp EEG. We propose the ERN from human EEG may be a particularly promising target for modeling using HNN’s elaborated neocortical network in future research.

### Limitations and future directions

The predictions in this study rely on the assumptions on the underlying HNN neocortical column model, which are based on generalizable features of neocortical circuitry and not specific to the potentially unique aspects of frontal cortical circuits. How much the finer details of microcircuit structure across brain areas contribute to macroscale EEG signals is unknown. Our results show that the FC-ERP shares similar features with sensory evoked ERPs from granular primary auditory and somatosensory areas (Jones et al., 2007; Kohl et al., 2022) and suggests that time-locked ERPs across the brain are constrained by conserved canonical structure of neocortical circuits and the laminar patterns of external inputs. HNN is designed to be a hypothesis testing and development tool using a pre-tuned, large-scale neocortical model. We have focused on testing specific *a priori* hypotheses in this study, with several alternatives examined, but we cannot rule out that additional dynamics may be involved in FC-ERP generation or stopping process. Overall, our results lead to many detailed predictions that provide targets for follow-up testing with invasive recordings (e.g., thalamic deep brain stimulation and recording, single-unit and laminar recordings in mPFC; Sherman et al., 2016) or other imaging (e.g., high-field fMRI, laminar MEG; Bonaiuto et al., 2021) modalities. HNN is developed with an open-source modular design so that if predictions are negated and/or new important circuit elements become known, it can be iteratively expanded and compared to data to account for new results. The HNN model tuned for (pre)frontal cortex dynamics presented here provides a unique starting point for further investigation of action-stopping and other cognitive control processes reflected in FC-ERPs in humans.

## Acknowledgements

This work was supported by funds from the NIH to DAD (T32MH126388), to JRW (R01 NS117753), and to SRJ (GR5271521, U24NS129945). The authors would like to thank Ryan Thorpe, Mainak Jas, Mohamed Sherif, and Maria Medalla for their helpful advice and feedback.

## Extended Data

**Figure 3-1.**
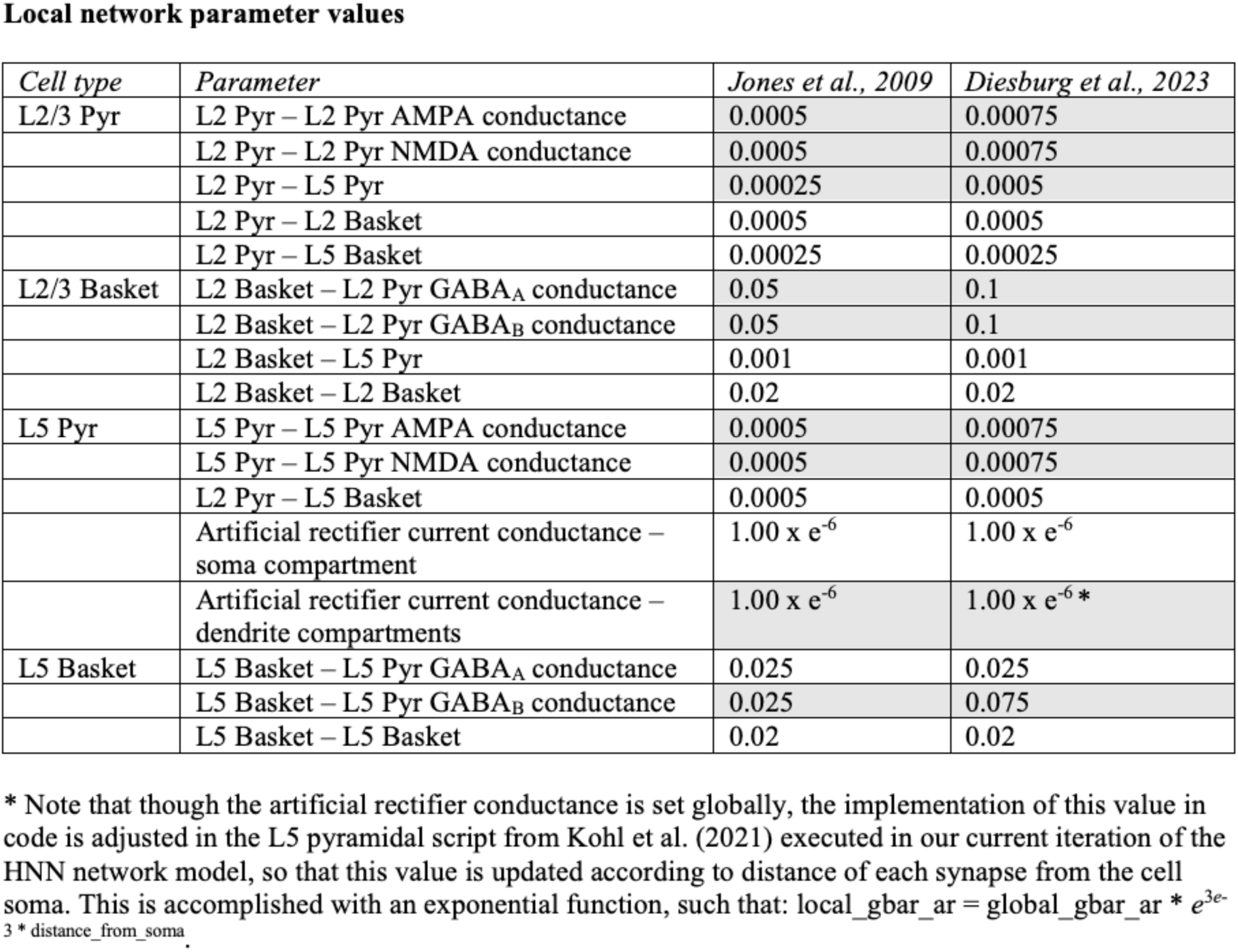
Changes in local network values implemented in the current model compared to the HNN default parameter settings from Jones et al. (2009).

**Figure 3-2.**
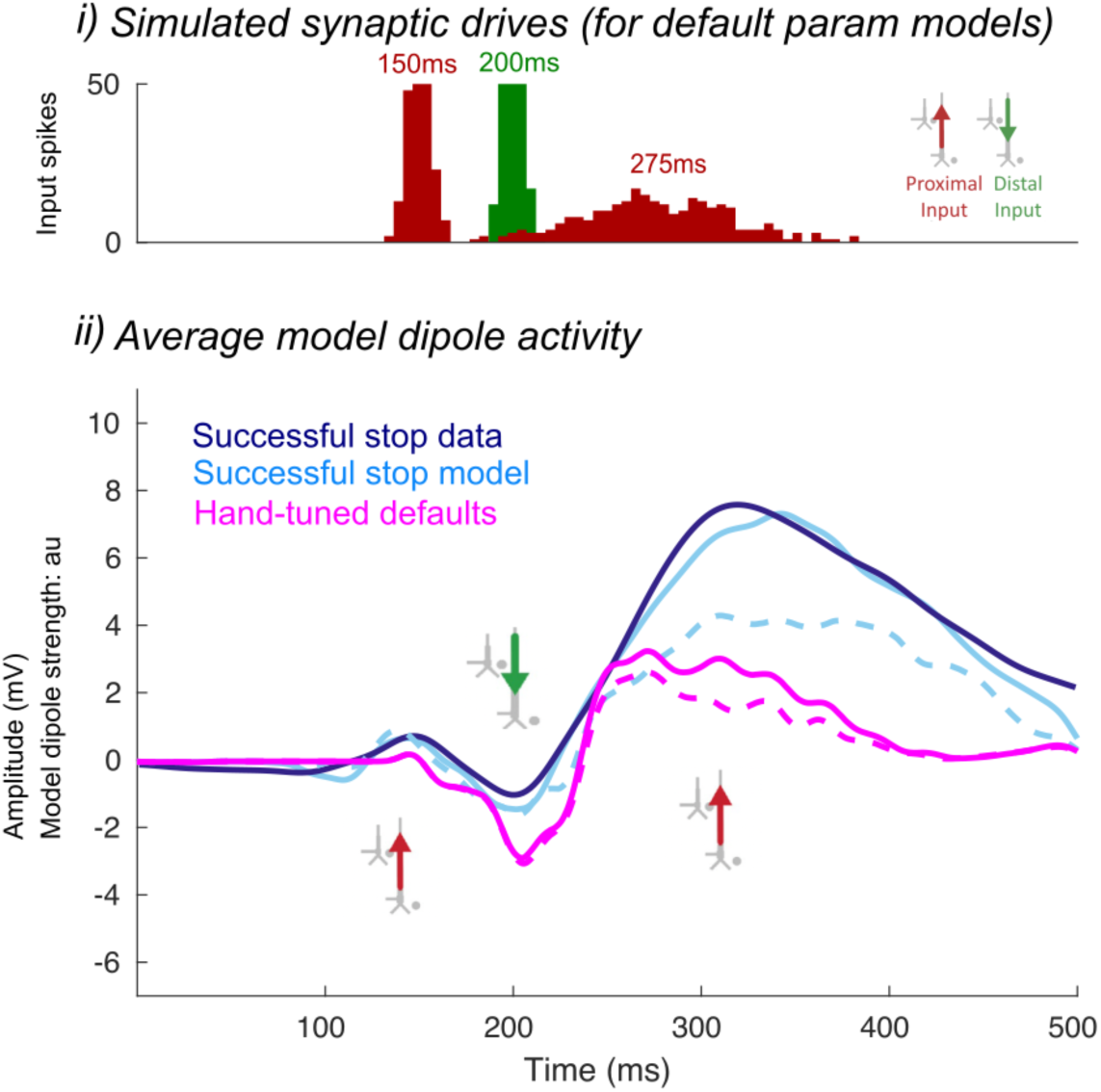
Preliminary HNN model for the successful Stop FC-ERP after rescaling and hand-tuning the mean onset and standard deviation of thalamocortical and cortico-cortical input drives. The dotted line shows model output with a re-emergent thalamocortical (Proximal 2) input of one spike and the solid line shows model output with a re-emergent thalamocortical (Proximal 2) of two spikes. Before adjusting the synaptic drive weights from the default settings (magenta), the amplitude difference in model output was not large. However, two spikes were needed in order to fit the FC-ERP because the difference in the amplitude between one-versus two-spike model outputs were larger once synaptic drive weights were adjusted (blue). i) All-trials histograms of thalamocortical and cortico-cortical drives delivered to the modeled column. ii) The average model dipole activity.

**Figure 3-3.**
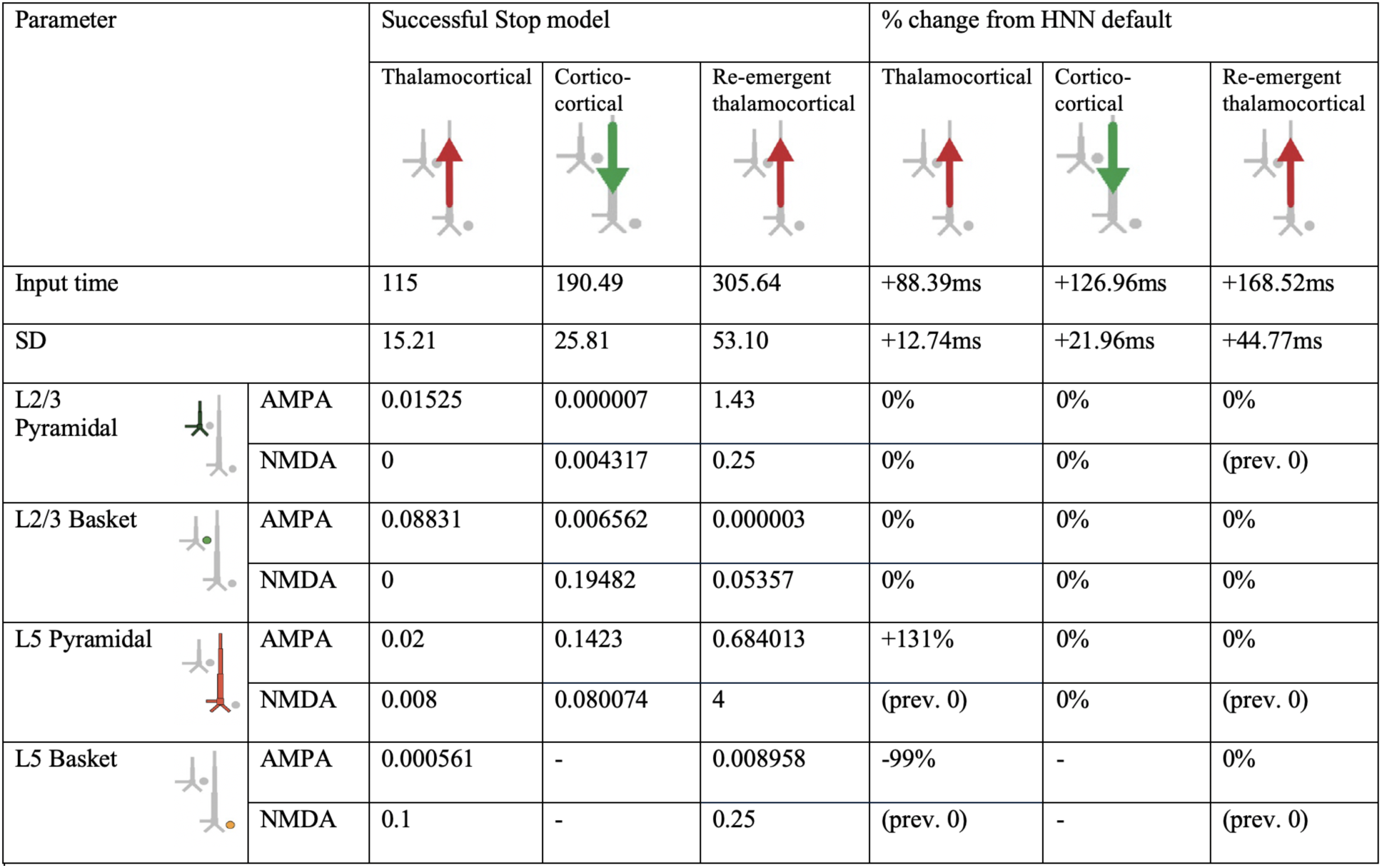
HNN model parameters for the successful Stop FC-ERP model (Figure 3A main text) and percentage changes from HNN default parameters after hand tuning and algorithmic optimization.

**Figure 3-4.**
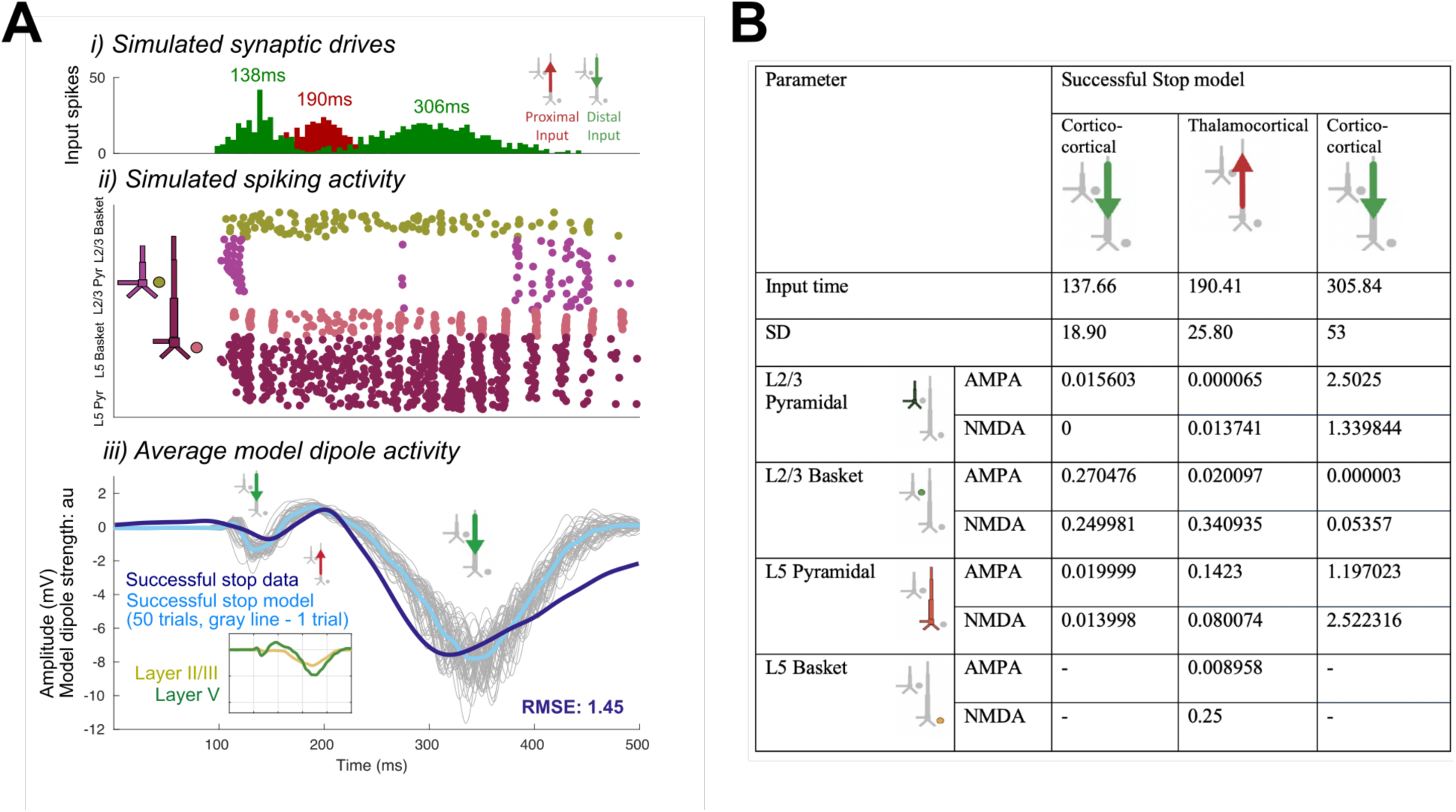
Optimized HNN models for the distal-drive-first alternate model of the successful Stop FC-ERP. A) Note that the second cortico-cortical (distal) drive does not produce dipole activity sustained for long enough during the P3 deflection to approximate the real ERP. i) All-trials histograms of thalamocortical and cortico-cortical drives delivered to the modeled column. ii) Simulated spikes from cell units of the model during one example trial. iii) The average model dipole activity and averaged dipoles in each model layer (thin gray lines indicate single-trial dipole activity, thick lines indicate the all-trial average). B) HNN model parameters for the model.

**Figure 3-5.**
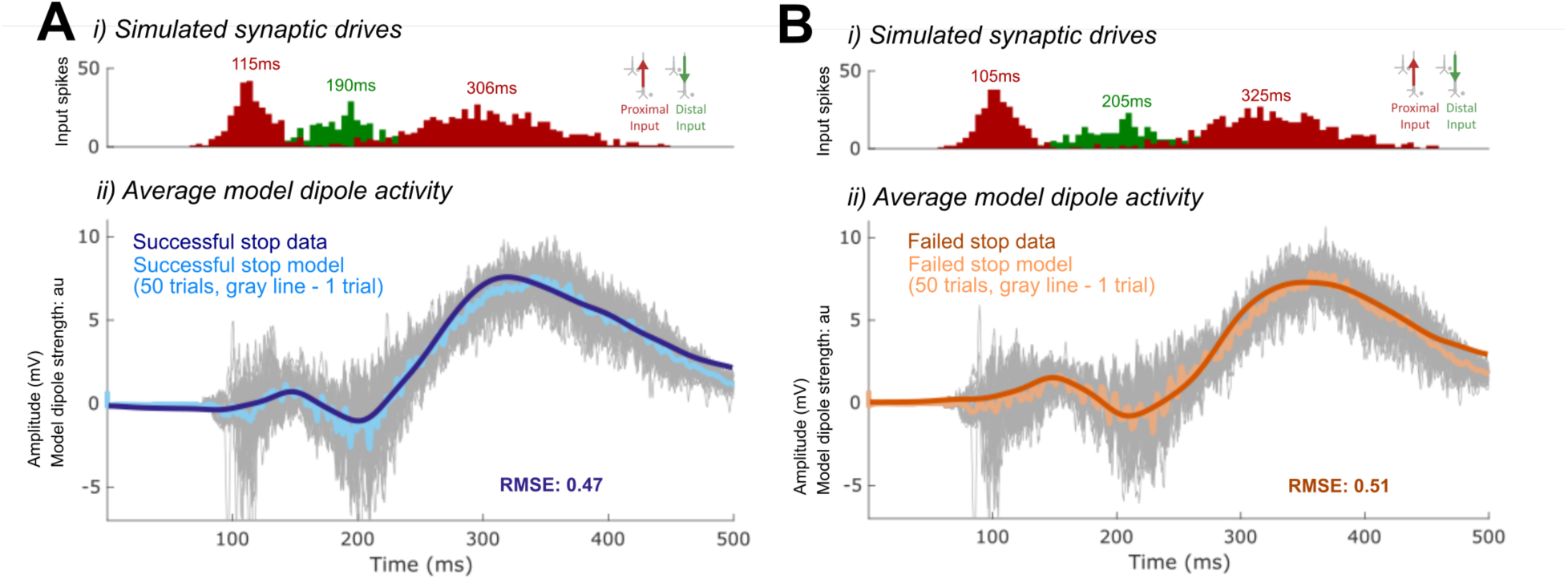
Examples of unsmoothed average dipoles from HNN models of successful (A) and failed Stop (B) trial evoked responses. i) All-trials histograms of thalamocortical and cortico-cortical drives delivered to the modeled column. ii) The average model dipole activity and averaged dipoles in each model layer (thin gray lines indicate single-trial dipole activity, thick lines indicate the all-trial average).

**Figure 3-6.**
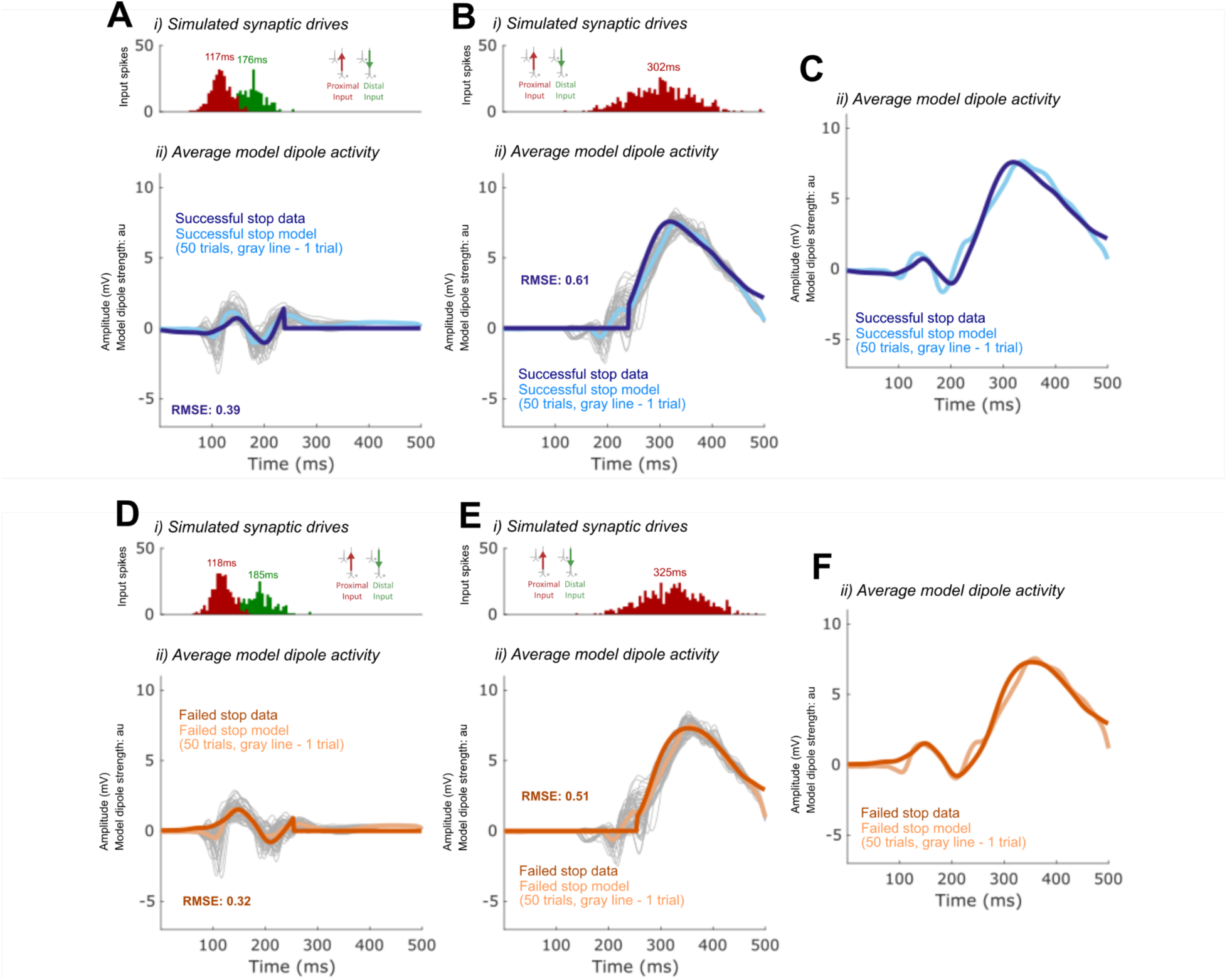
Predicting mechanisms of the FC-ERP using two modeled sources (i.e., neocortical columns) instead of one. A) Parameters associated with the first proximal and distal drive in the successful Stop model in Fig. 3A were optimized to fit only the P2 and N2 deflection of the ERP. The later parts of the waveform are ignored and set to zero. B) Parameters associated with the second proximal drive in the successful Stop model in Fig. 3A were optimized to fit only the P3 deflection of the ERP. The earlier parts of the waveform are ignored and set to zero. C) The average dipole output from panels A and B added together, plotted alongside the empirical successful Stop FC-ERP. D) Parameters from panel A were optimized to fit the P2 and N2 deflection of the failed Stop FC-ERP. The later parts of the waveform are ignored and set to zero. E) Parameters from panel B were optimized to fit the P3 deflection of the failed Stop FC-ERP, with only timing and deviation of the drives allowed to vary. The earlier parts of the waveform were ignored and set to zero. F) The average dipole output from panels A and B added together, plotted alongside the failed successful Stop FC-ERP. In all panels, i) all-trials (n=50) histograms of thalamocortical and cortico-cortical drives delivered to the modeled column. ii) the average model dipole activity and averaged dipoles in each model (thin gray lines indicate single-trial dipole activity, thick lines indicate the all-trial average).

**Figure 4-1.**
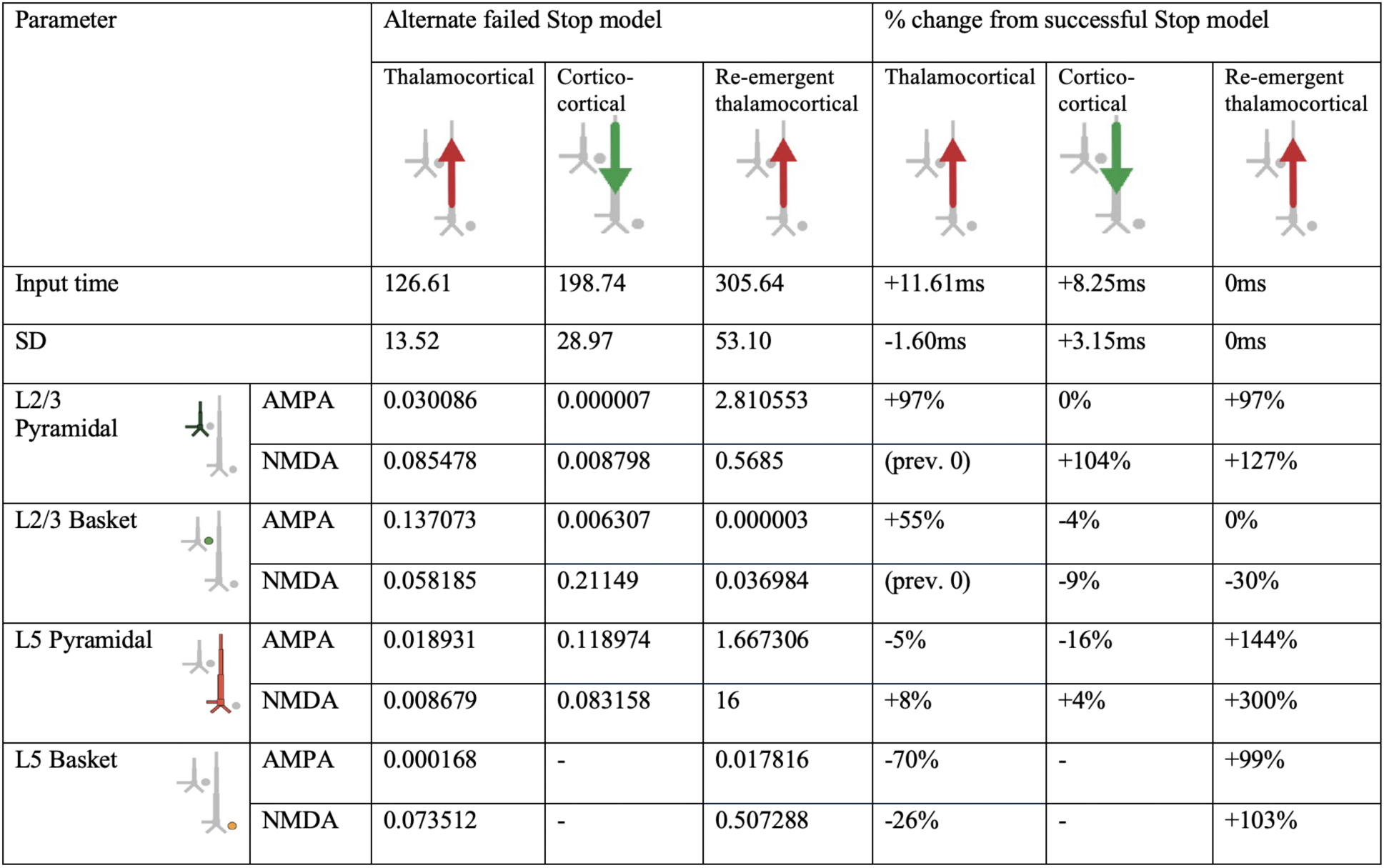
HNN model parameters for an alternative model of the failed Stop FC-ERP model in which one spike was used for the second proximal drive rather than two (Figure 4A main text). A round of optimization within 300% parameter change was run starting with the successful Stop parameters (with one spike rather than 2 for the re-emergent thalamocortical Proximal 2 drive).

**Figure 4-2.**
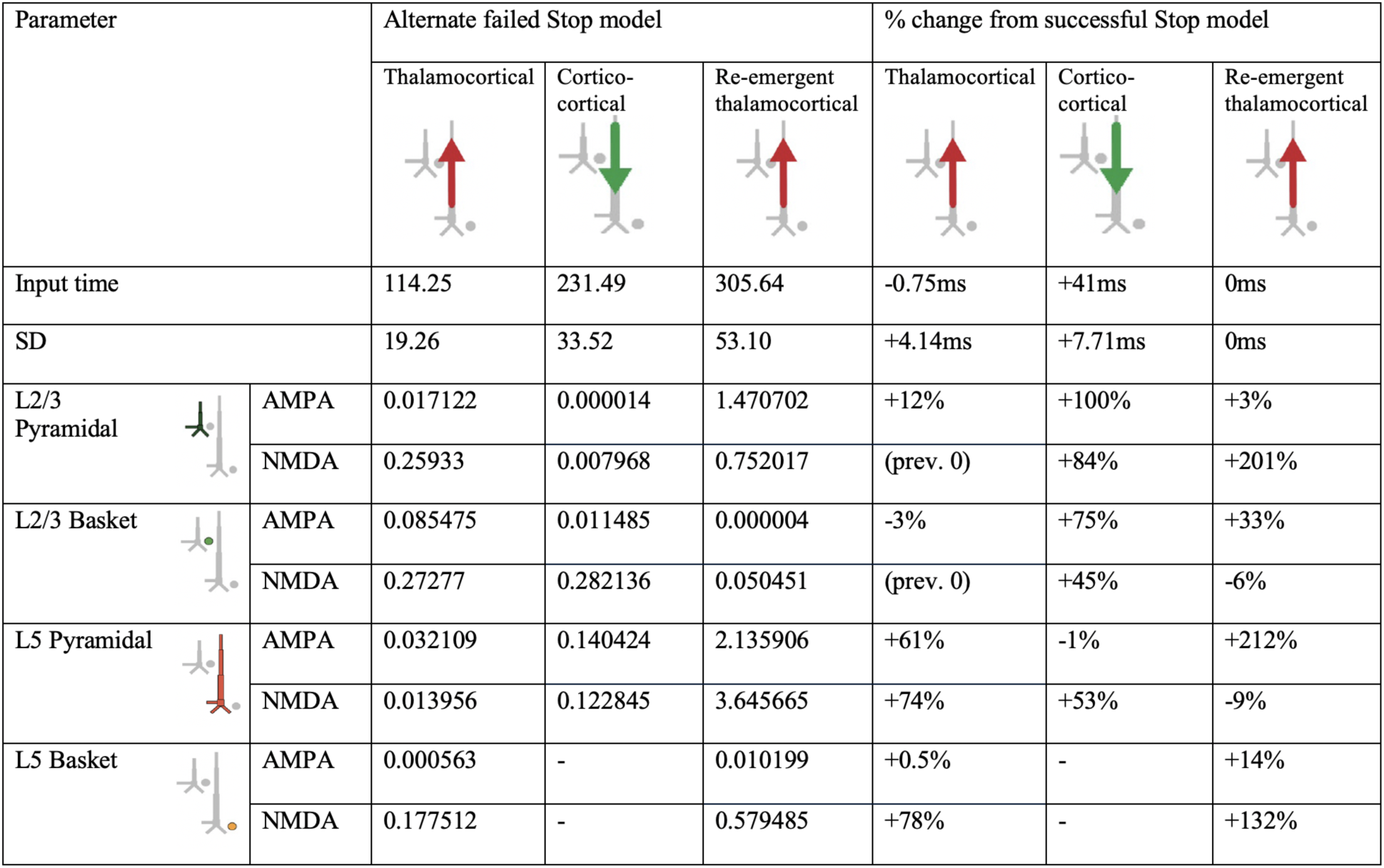
HNN model parameters for an alternative model of the failed Stop FC-ERP model in which synaptic weights but not timing of the second proximal drive were allowed to vary from the successful Stop model (Figure 4B main text).

**Figure 5-1.**
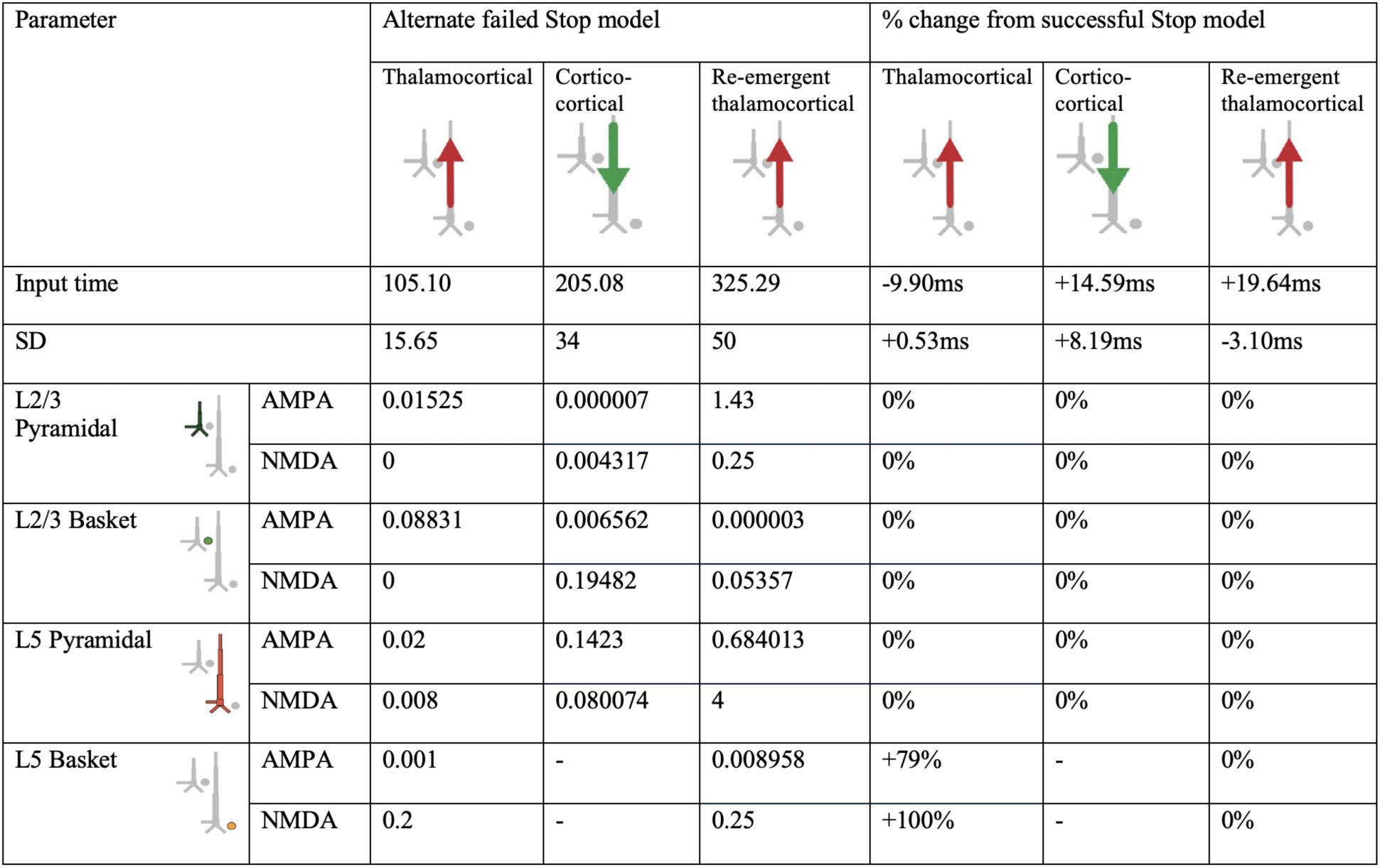
HNN model parameters for an alternative model of the failed Stop FC-ERP model in which synaptic weights in Layer 5 during the distal drive were reverted to successful Stop model weights and Layer 5 basket cell weights during the first proximal drive increased (to simulate somatic inhibition; Figure 5A main text).

**Figure 5-2.**
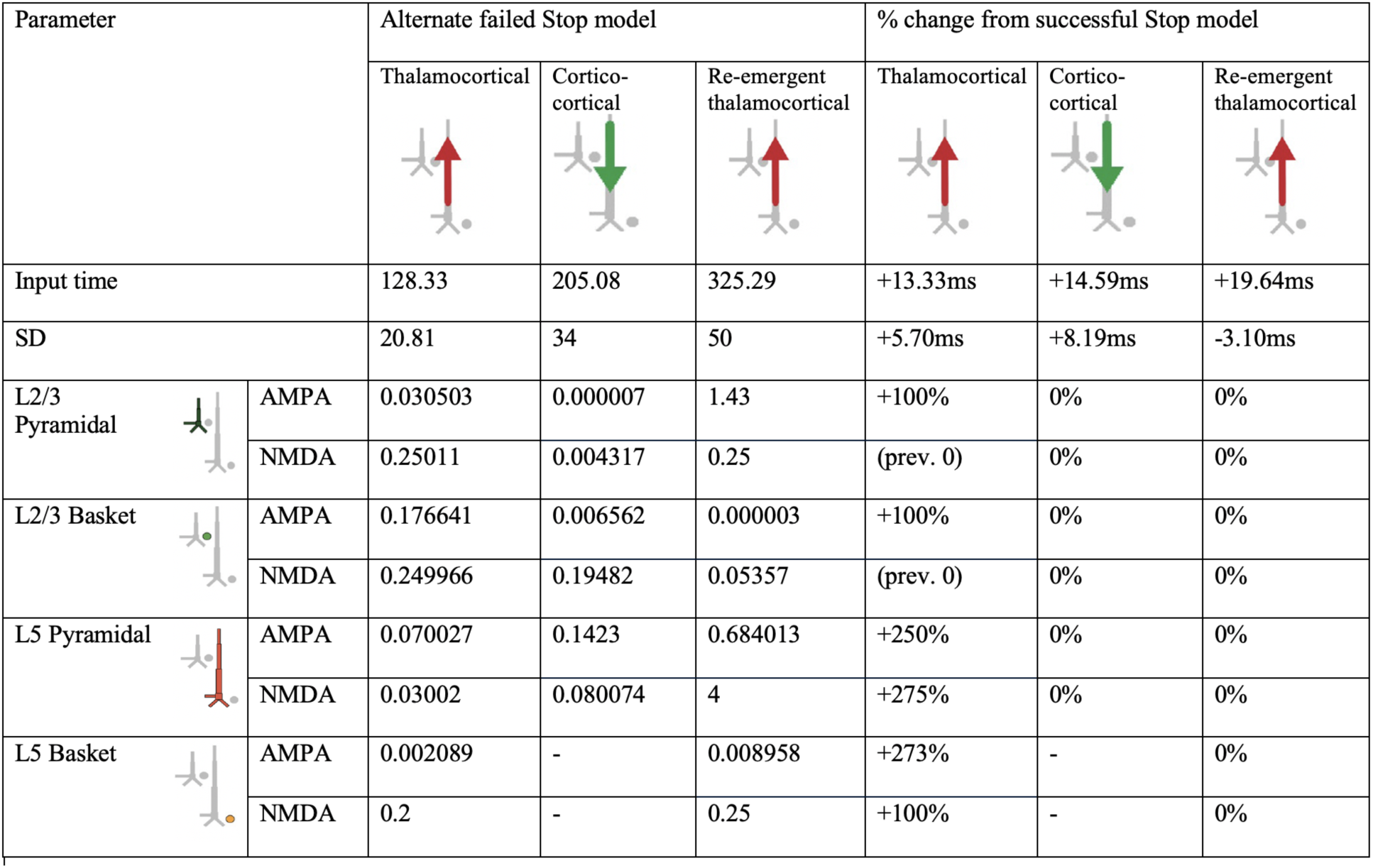
HNN model parameters for an alternative model of the failed Stop FC-ERP model in which all synaptic weights of Proximal 1 drive were optimized beginning from the somatic inhibition condition simulated in the alternate model presented in Figure 5A (Figure 5B main text). The timing and deviation of the Proximal 1 drive was similarly optimized.

